# Neuronal identity defines α-synuclein and tau toxicity

**DOI:** 10.1101/2022.06.24.496376

**Authors:** Roman Praschberger, Sabine Kuenen, Nils Schoovaerts, Natalie Kaempf, Jasper Janssens, Jef Swerts, Eliana Nachman, Carles Calatayud, Stein Aerts, Suresh Poovathingal, Patrik Verstreken

## Abstract

Pathogenic α-synuclein and tau are critical drivers of neurodegeneration and their mutations cause neuronal loss in patients. Whether the underlying preferential neuronal vulnerability is a cell-type intrinsic property or a consequence of increased expression levels is an open question. Here, we explore cell-type specific α-synuclein and tau expression in human brain datasets and use deep phenotyping as well as brain-wide single-cell RNA sequencing of >200 live neuron types in fruit flies to ask which cellular environments react most to α-synuclein or tau toxicity. We detect phenotypic and transcriptomic evidence of differential neuronal vulnerability independent of α-synuclein or tau expression levels. Comparing vulnerable with resilient neurons enabled us to identify molecular signatures associated with these differential responses. We used these to verify, and then predict resilient and vulnerable neuron subtypes in human brains. This confirms *substantia nigra* dopaminergic neurons to be sensitive to α-synuclein, and we predict pathogenic tau vulnerable and protected cortical neuron subtypes. Our work indicates that cellular determinants confer selective vulnerability to specific types of amyloid toxicity, thus paving the way to leverage neuronal identity to uncover modifiers of neurodegeneration-associated toxic proteins.

## Introduction

Intracellular α-synuclein or tau aggregates are neuropathological hallmarks of several neurodegenerative diseases, including Parkinson’s and Alzheimer’s disease. They are characterized by selective neuronal vulnerability, where some neurons are more affected than others (Fu et al., 2018). Since in rare familial forms α-synuclein and tau mutations are also the primary cause for disease (Hutton et al., 1998; Polymeropoulos et al., 1997), they represent a clear-cut way to assess whether the identity of neuronal cell type can modulate α-synuclein or tau toxicity.

α-synuclein mutations lead to Parkinson’s disease (Goedert et al., 2013). The core affected brain regions are the *substantia nigra, locus coeruleus*, temporal cortex and hippocampus (Schneider and Alcalay, 2017). Tau mutations typically give rise to a form of frontotemporal dementia with parkinsonism where the main affected brain regions are the frontal and temporal cortices and certain subcortical and brain-stem areas (Mirra et al., 1999; Spillantini et al., 1998; Tsuboi, 2006). Thus, neuropathology and clinical signs indicate that also α-synuclein and tau mutations are associated with region and cell-type specific effects, which are likely exerted through a toxic gain-of-function. This is suggested by the dominant inheritance of the mutations, their aggregation promoting effects and induction of neurodegeneration when expressed in different mammalian and invertebrate *in vivo* models (Feany and Bender, 2000; Flagmeier et al., 2016; Nacharaju et al., 1999; Van Der Putten et al., 2000; Wittmann et al., 2001; Yoshiyama et al., 2007).

α-synuclein and tau are expressed brain-wide (Lein et al., 2007; Lizio et al., 2015, 2019), but it is not known whether preferential regional and cell-type vulnerability in mutation carriers is the result of an interaction of these toxic proteins with particular features of different neuron types, or if it is the simple consequence of increased expression levels in vulnerable neurons. Rigorously addressing this question is important, since if it were to be established that some neurons are preferentially affected beyond the contributions of increased expression levels, it provides the basis for unravelling the mechanisms of neuronal resilience in the face of α-synuclein or tau proteopathic stress.

Here we use public human single-cell gene expression atlases to show that α-synuclein and tau are highly expressed in vulnerable *and* resilient neurons. We experimentally confirm that α-synuclein and tau mutations have the ‘intrinsic potential’ to differentially affect neurons by combining neuronal phenotyping and brain-wide single-cell RNA sequencing in novel, highly controlled *Drosophila* models. This approach enables us to simultaneously assess the transcriptional deregulation in >200 diverse conditions (neuron subtypes) when they express α-synuclein or tau, compare the contributions of transgene expression levels and perform cell-type specific experimental validation. This reveals differential vulnerability to α-synuclein and tau proteopathic stress independent of expression levels. Resilient neurons distinguish from vulnerable neurons by their differential expression of mitochondrial respiration in the case of α-synuclein and axon/synapse organization when expressing tau. We finally use vulnerability/resilience defining genes in *Drosophila* to validate and then predict vulnerable/resilient cell types in the human brain. Our work represents an important step in leveraging neuronal diversity to uncover α-synuclein and tau toxicity-modifying pathways and genes.

## Results

### Cell-type specific α-synuclein and tau expression levels do not account for differential vulnerability in mutation carriers

To test whether increased expression of α-synuclein or tau explains cell-type preferential vulnerability, we explored publicly available human brain mRNA sequencing datasets of non-neurodegeneration control individuals. In regional bulk gene expression data across different human brain areas, there is brain-wide expression of α-synuclein and tau without obvious correlation with regions known to be differentially vulnerable (Figure S1A and S1B). Expression of α-synuclein and tau is higher in the cerebellum as compared to the primary sites of vulnerability, i.e. *substantia nigra* and frontal cortex, respectively (Figure S1A and S1B). Median α-synuclein and tau expression in the hippocampus and *substantia nigra* – both regions affected in mutations carriers (Mirra et al., 1999; Schneider and Alcalay, 2017; Spillantini et al., 1998) – are amongst the 5 lowest expressing brain regions out of 13 analyzed. Hence, α-synuclein or tau expression levels in specific brain regions do not correlate with known vulnerable regions.

Next, to assess the contribution of individual cell types to regional bulk gene expression, we analyzed α-synuclein and tau in single-cell RNA sequencing (scRNA-seq) datasets. We resorted to the nervous system-wide atlas of mouse (Zeisel et al., 2018), that allows us to compare relative expression levels across most brain cell types. Patients with pathogenic α-synuclein, and frequently also those harboring tau mutations, show loss of melanin-containing dopaminergic neurons (Mirra et al., 1999; Schneider and Alcalay, 2017; Spillantini et al., 1998). In the mouse brain, median α-synuclein and tau expression in *substantia nigra* and ventral tegmental area dopaminergic neurons are within the top decile of cell types (Figure S1C and S1D). Furthermore, median α-synuclein expression in CA3 hippocampal neurons that substantially degenerate in α-synuclein mutation carriers (Spira et al., 2001), ranks high (5^th^) across 265 cell types (Figure S1C). However, also many other cell types not reported as affected in α-synuclein and tau mutation carriers express high levels of α-synuclein and tau, such as spinal cord excitatory neurons (Figure S1C) or cranial nerve nuclei (Figure S1D). Collectively, this suggests that high α-synuclein or tau expression levels might sensitize neurons, but that other cellular features exist that modulate toxicity.

We confirm these findings using single-cell sequencing datasets that are available from specific human brain regions. To enable comparisons between datasets acquired with different technologies, we further normalized mean α-synuclein and tau expression to the means of all detected genes in a given cell type by calculating their expression percentiles. In dopaminergic neurons in a combined *substantia nigra* and middle frontal gyrus dataset, we find robust expression of α-synuclein and tau, both above the 97^th^ percentile (Agarwal et al., 2020) (Figure S1E-S1H). Also, the preferentially vulnerable CA2 and CA3 neurons express high levels of α-synuclein in the human hippocampus (Franjic et al., 2022) (Figure S1I and S1J). Tau expression is similarly high across different neuron types of the entorhinal cortex, subiculum and hippocampus (Figure S1K and S1L). Across multiple human cortical areas α-synuclein expression is higher in excitatory compared to inhibitory neurons (Figure S1M and S1N). Tau is expressed highly throughout excitatory and inhibitory neurons and expression levels are similar across brain areas, providing no direct explanation why frontal and temporal cortices are preferentially vulnerable in tau mutant patients (Figure S1O and S1P).

Taken together, our analysis of regional and cell-type specific α-synuclein and tau expression levels across the human and mouse brain reveals that vulnerable cell types and brain regions express high levels of α-synuclein and tau. However, similar or sometimes even higher levels are observed in several regions and cell types that are not apparently vulnerable in α-synuclein and tau mutant patients, suggesting that region and cell-type intrinsic properties influence toxicity.

### *Drosophila* models of α-synuclein and tau mutations show differential neuronal vulnerability

To experimentally test whether neurons are differentially affected when exposed to pathogenic α-synuclein and tau, we generated *Drosophila* models expressing human A53T mutant α-synuclein or P301L mutant tau in all neurons (Figure 1A). In fruit flies, a very large number of highly diverse neuron types can be assayed in parallel (Davie et al., 2018; Jenett et al., 2012; Konstantinides et al., 2018; Kurmangaliyev et al., 2020; Li et al., 2022; Özel et al., 2021). We also generated, as extra controls, transgenic flies with an empty backbone insertion (mini-*w*^*+*^), a fluorophore-dead GFP (smGdP) and α-synuclein as well as tau mutants where the main aggregation-prone regions are deleted (NAC and PHF6 VQIVYK, respectively; von Bergen et al., 2000, 2001; Bodles et al., 2001; Rodriguez et al., 2015; Sawaya et al., 2007). These were all crossed for >5 generations into the same inbred reference background and expressed using the neuronal *nSyb* promoter, which acts through the exogenous QF2w transcriptional activator and QUAS binding sites (Figure 1A) (Potter et al., 2010; Riabinina et al., 2015). Two design features are of particular importance. First, because all transgenes are inserted as single copies into the same genomic locus (Bischof et al., 2007; Venken et al., 2006) (Figure 1B), their relative expression levels across cell types is genetically hardwired to be comparable – which we confirm with single-cell RNA sequencing (see section below). Second, α-synuclein and tau are expressed at levels comparable to those found in human brains (Figure 1C-1F). We also validated that the expression is maintained throughout the lifespan of these flies (Figure S2A-S2D), enabling us to include ageing as an experimental variable.

**Figure 1.**
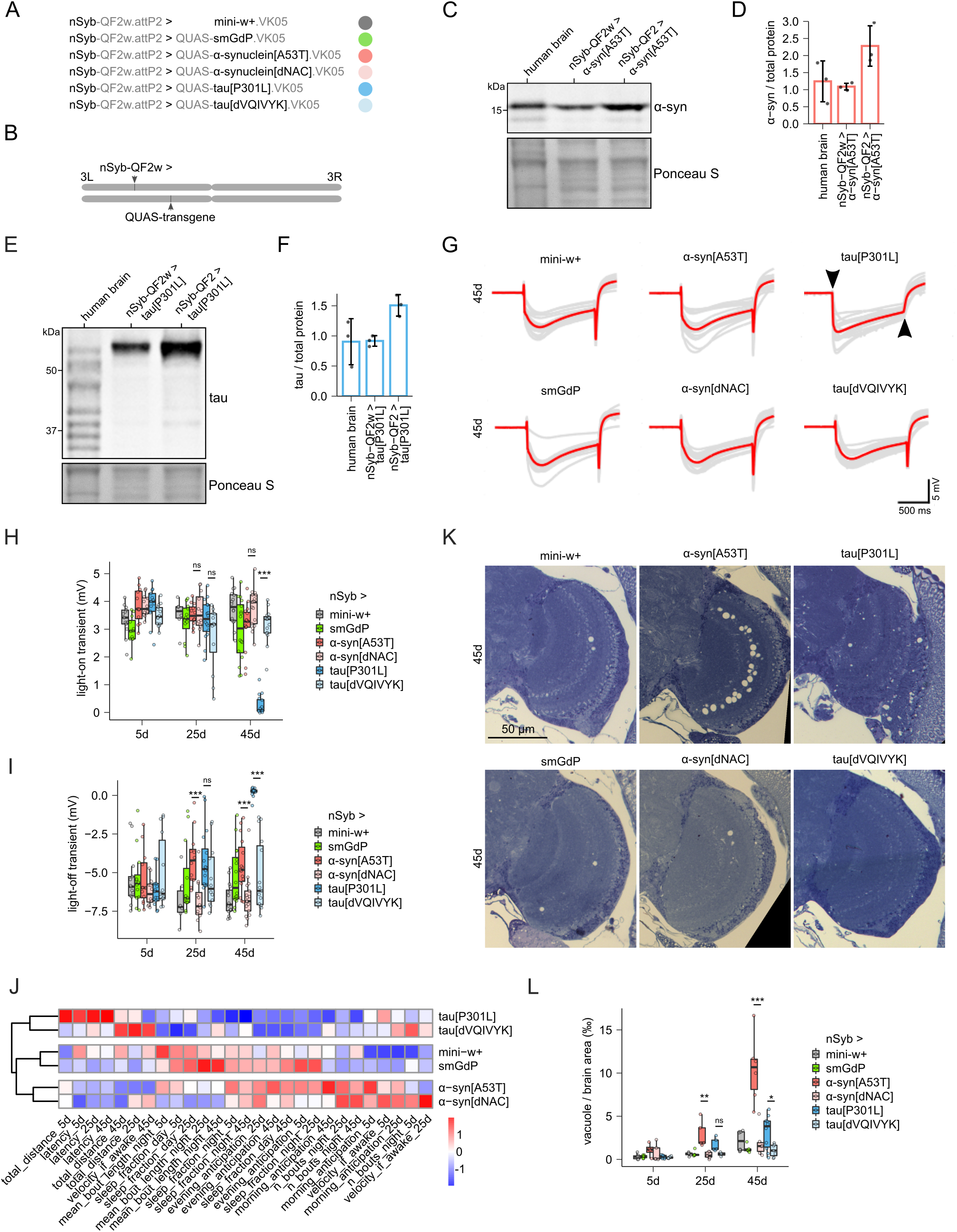
Divergent *Drosophila* phenotypes as a result of differential neuronal vulnerability to pathogenic α-synuclein and tau. (A) Full genotypes of the experimental lines plus associated controls generated in this study. (B) Schematic of the model genetics. Transgenes are inserted as single copies in the same insertion site on chromosome 3. They are expressed across all neurons upon a single cross with the pan-neuronal synaptobrevin (*nSyb*) promoter. (C-F) Representative immuno-blots and quantification of α-synuclein and tau expression in fly heads as compared to healthy human brains. We use the weaker expressing QF2w (middle lanes) as compared to the stronger QF2 (Riabinina et al., 2015). Of note, we are expressing only one tau splice isoform (0N4R) in our *Drosophila* models, which is contrasted by multiple tau bands in human brains. Replicate values, mean and standard deviation (SD) are shown. (G) ERG traces of repeat experiments (gray) and mean (red) for 45 day old flies are shown. Arrowheads indicate the loss of light-on and off transients in P301L tau expressing flies. (H, I) Quantification of ERG light-on/-off transients at 5, 25 and 45 days. Replicate values, median, IQR and whiskers at 1.5xIQR are shown. ns, p > 0.05; ***p < 0.001; Wilcoxon rank-sum test and Bonferroni correction. (J) Hierarchical clustering across scaled mean sleep and activity parameters at 5, 25 and 45 days of age. Z-scores of condition means are shown. N = 53-89 flies per genotype and age; 3 independent experiments each. (K) Representative microscope images of toluidine blue stained brain section at age 45 days. Optic lobes and part of adjacent central complexes are shown. (L) Automated quantification of vacuole area across entire brains normalized for brain size. Replicate values, median, IQR and whiskers at 1.5xIQR are shown. ns, p > 0.05; *p < 0.05; **p < 0.01; ***p < 0.001; Two-sample t-test and Bonferroni correction.

To test the functional impact of pathogenic α-synuclein and tau in a well-established model circuit, we performed electro-retinogram (ERGs) recordings (Figure 1G-1I and Figure S2E and S2F). ERGs are local field potentials that occur in response to a light stimulus, thus providing the integrated response of the retinal cells and post-synaptic lamina monopolar neurons. At 45 days of age, but not before, we find in P301L tau expressing flies a complete loss of light-on and -off transients that are physiologically elicited at the switchover from dark to light and vice versa (Figure 1G-1I and Figure S2E). Conversely, animals expressing tau harboring an aggregation-domain deletion do not show ERG defects, indicating this is largely driven by aggregation toxicity. ERGs in A53T α-synuclein expressing flies and the aggregation-domain deletion are similar to controls (Figure 1G-1I and Figure S2E). Given the difference between ERG recordings between α-synuclein and tau expressing flies, these data are functional evidence of differential neuronal vulnerability in this specific circuit.

To evaluate the functional integrity of broader neuronal circuits upon expression of the toxic proteins, we used high-resolution sleep and activity monitoring using video-tracking of individual flies (Geissmann et al., 2017) and extracted sleep and activity parameters that are controlled by different neuronal systems systems (Donlea et al., 2011; Valadas et al., 2018). Mutant α-synuclein and tau flies cluster on opposing ends of the hierarchical tree of behavioral parameters at 5, 25 and 45 days of age (Figure 1J), consistent with different circuits being affected. Furthermore, the distance between α-synuclein/tau harboring aggregation-domain deletions vs. their pathogenic counterparts is larger as compared to the distance between the mini-*w*^*+*^ and smGdP controls. This suggests that the observed circuit dysfunction is partly caused by aberrant aggregation mediated toxicity.

Finally, we also tested whether pathogenic α-synuclein and tau cause neurodegeneration in our models. We generated brain sections and used automated analysis to quantify the area of brain vacuolization (Figure 1K and 1L). A53T α-synuclein-expressing brains show numerous vacuoles in the optic lobes at 25 days of age (Figure 1L). This is even more apparent at 45 days of age and is not observed in α-synuclein with the aggregation-domain deletion. P301L tau expressing flies also exhibit age-dependent increases of vacuolar area that is not seen in animals expressing tau harboring an aggregation-domain deletion (Figure 1L). However, the extent of vacuolization and the regional distribution is markedly different to our observations in A53T α-synuclein-expressing flies. In P301L brains vacuoles are present throughout the brain; in A53T α-synuclein they mainly cluster in the optic lobes (Figure 1K). This further suggests that different neurons are affected in A53T α-synuclein vs. P301L tau brains.

### Differential transcriptomic impact to pathogenic α-synuclein and tau across >200 neuron types

To identify neuron types differentially vulnerable to α-synuclein and tau we performed brain-wide single-cell RNA sequencing. We sampled A53T α-synuclein, P301L tau, mini-*w*^*+*^ controls and smGdP at 5 and 25 days of age in six repeat experiments (Figure 2A). Cells from all genotypes and experimental dates intermix homogenously, suggesting that all cell types are formed across conditions (Figure 2A and Figure S3A-C). After quality control filtering, we retain 143013 cells, evenly distributed between the conditions (Figure S3D and S3E). Individual cells were clustered with the Leiden community detection algorithm and annotated with classifiers previously trained on manually annotated fly brain atlases as well as with bulk transcriptomes of FAC-sorted cell types (Figure 2B) (Davis et al., 2020; Li et al., 2022; Özel et al., 2021; Traag et al., 2019; Pech et al., unpublished). We identify a total of 208 brain cell types of which 202 are neurons of all the major neurotransmitters classes (Figure S4A-S4D). This large number of neuronal subtypes is the consequence of being able to sample entire fly brains in every repeat experiment. Our dataset thus provides a high-resolution platform for the study of the neuron-type specific impact of pathogenic α-synuclein and tau.

**Figure 2.**
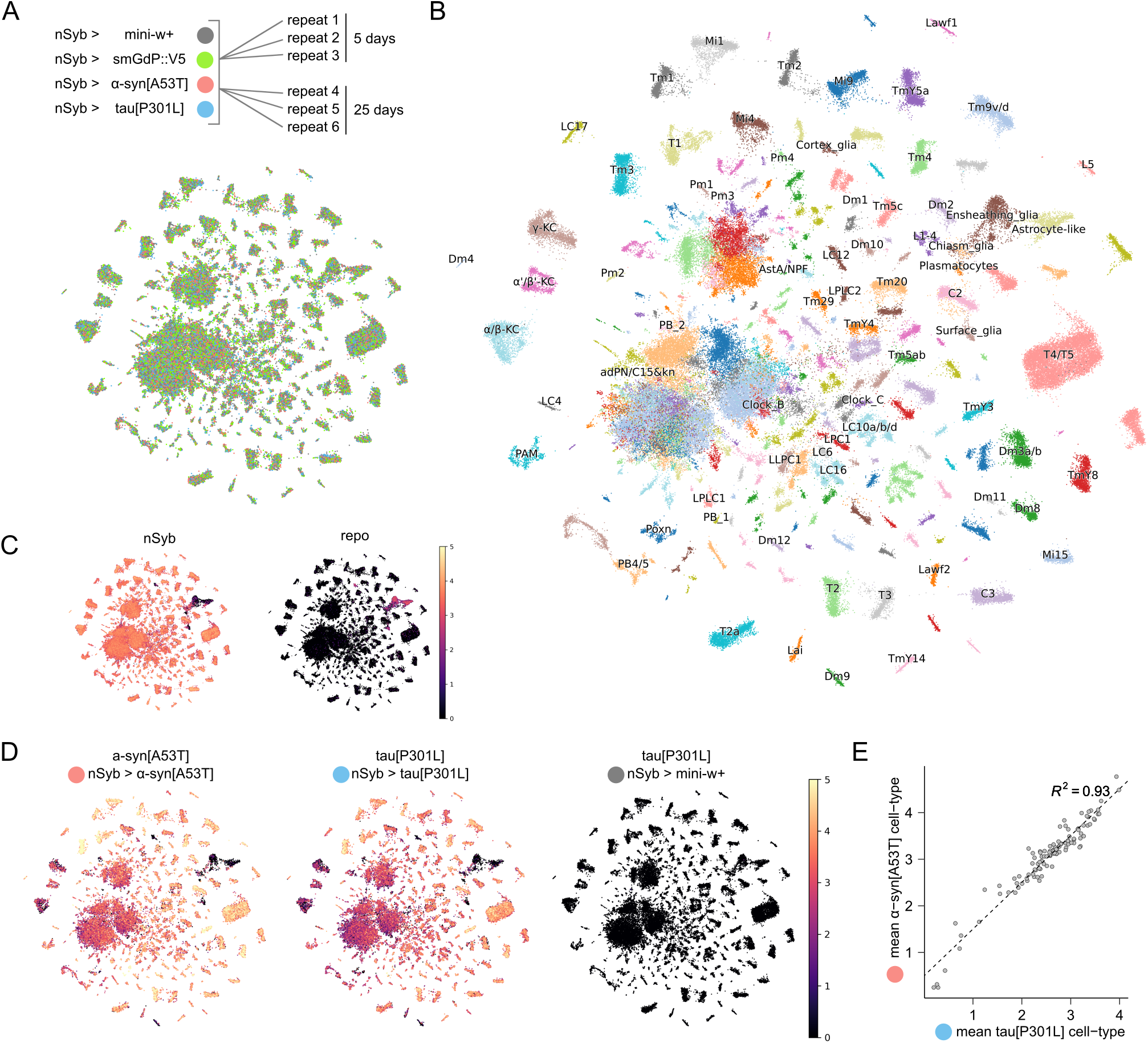
A high-resolution atlas of α-synuclein and tauopathy *Drosophila* model brains. (A) Experimental design of this study. All three genotypes were processed in parallel. UMAP depicting even intermixing of genotypes without the use of integration algorithms. (B) UMAP with cluster labels is shown. In total 67 cell types could be identified. For the rest, we inferred their neuronal identity as well as neurotransmitters based on marker genes (not shown). (C) Normalized and log-transformed *nSyb* (neuronal marker) and *repo* (glial marker) expression levels across all cells are shown. (D) Normalized and log-transformed A53T α-synuclein and P301L tau expression levels across cells from each of the three genotypes is shown. (E) Scatter-plot and linear regression of mean normalized and log-transformed transgene levels across cell types of 5 day and 25 day old P301L tau vs. A53T α-synuclein models. Only cell types with a minimum of 50 cells and those that are not part of the large central clusters are shown.

We first confirmed that A53T α-synuclein and P301L tau are expressed at similar levels in the identified cell types; a prerequisite to study their differential impact. Consistent with the experimental design, we detect A53T α-synuclein and P301L tau in *nSyb*-positive cells (Figure 2C), and we observe highly correlated expression in all analyzed cell types (Figure 2D-E and Figure S4E and S4F). Conversely glial cells, which express the *repo* marker gene, or mini-*w*^*+*^ do not express the toxic transgenes (Figure 2C-D). We also see that cell-type specific expression levels are maintained throughout ageing from 5 to 25 days (Figure S4G and S4H), suggesting that comparing cell-type specific effects at 5 and 25 days are not confounded by changes in transgene levels.

Next, we calculated gene expression changes in cells of A53T α-synuclein, P301L tau and smGdP compared to non-expressing controls for each cell type. We filtered cell types with low cell counts as well as poorly defined clusters and evaluated differentially expressed genes (DEGs) both at 5 and 25 days separately as well as in a combined model with DESeq2 (Love et al., 2014) (Figure 3A-3B and Figure S5). The magnitude of transcriptional deregulation across neuron types is highly heterogenous, ranging from 1 DEG in glutamatergic neuron type 44 (N-44-Glut) and LC12 neurons to 36 in T1 neurons upon expression of A53T α-synuclein in the 5 and 25 day joint analysis; and from 1 DEG in the GABAergic neuron type 75 (N-75-GABA) to 78 in T1 neurons when P301L tau is expressed (Figure 3A and 3B). Thus, some neuron types are deregulated several-fold stronger than others. Similarly, another well-established differential gene expression testing algorithm, edgeR (Robinson et al., 2009), provides robustly correlated prioritizations of neuron types based on magnitudes of transcriptional deregulation as compared with DESeq2 (median Spearman rho = 0.655 at the 5, 25, and 5 + 25 days analyses). To further refine our neuron type prioritizations based on transcriptomics, we evaluated technical covariates that might influence the number of recovered DEGs. The number of cells per neuron type shows a moderate contribution (Spearman rho < 0.5, Figure S6A and S6B); the number of detected genes is not correlated in A53T α-synuclein and only weakly correlated in P301L tau due to a few outliers (Spearman rho < 0.3, Figure S6C and S6D), such as T1 neurons which are physically larger than many other fly brain neuron types. Correspondingly, in T1 neurons ∼70% more genes are detected. Finally, we also assessed whether transgene levels predicted the number of DEGs and found that A53T α-synuclein does not correlate and P301L tau moderately (Spearman rho = 0.13 and 0.43, Figure S6E and S6F). To correct for the technical covariates, we fitted negative binomial regression models including these explanatory variables. We also included toxic transgene levels, which is critical in order to be able to assess neuron-type specific effects *beyond* and independent of expression levels. This ultimately enabled us to make conservative estimates as to which neuron types exhibit significantly more or less DEGs than predicted by the model (Figure S6G and S6H). We find that TmY5a, Tm3, α/β-KC, TmY8, T2a, Tm1 neurons are impacted significantly stronger than expected upon A53T α-synuclein expression; likewise, Tm3, C2, T2, C3, Tm1, T3 and Mi15 upon P301L tau expression (Figure S6G and S6H), independent of the covariates cell and gene number as well as transgene levels. While Tm3 and Tm1 neurons appear preferentially deregulated both upon A53T α-synuclein and P301L tau expression, there are several neuron types where the responses in the A53T α-synuclein *versus* P301L tau *versus* smGdP conditions diverge, such as C2, C3, T2, T3, Tm2, α/β-KC and TmY5a neurons (Figure S6I). Collectively, these results provide transcriptomic evidence of differential neuron impact by α-synuclein and tau.

**Figure 3.**
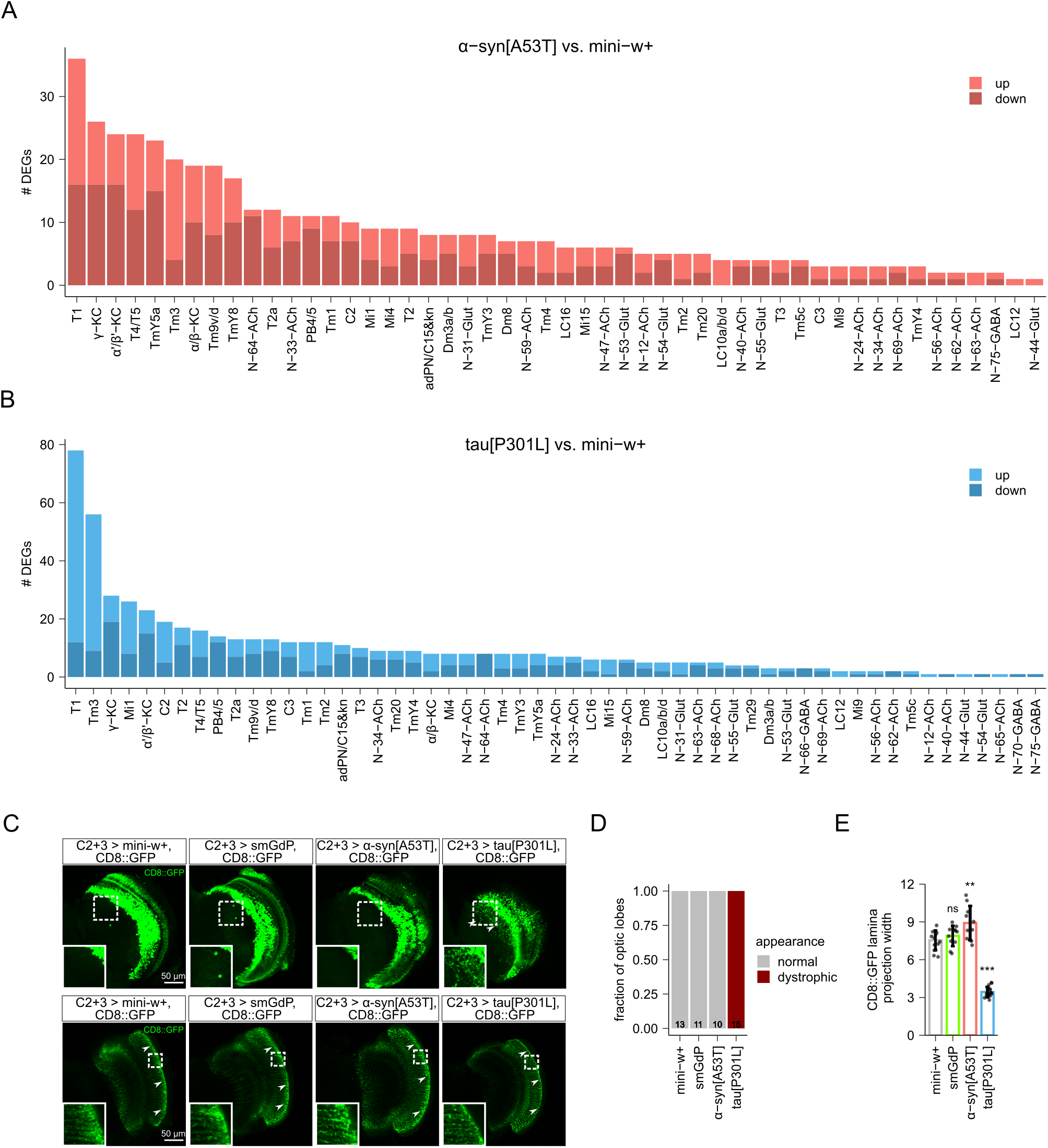
Neuron-type specific transcriptional and structural impact in α-synuclein and tauopathy *Drosophila* models. (A, B) The number of significantly up-/downregulated DEGs per neuron type are shown for α-synuclein[A53T] and tau[P301L] vs. mini-w+. Only neuron types that have >10 cells per repeat, >130 total mutant cells and do not map to poorly defined clusters are shown. DEG testing was performed with DESeq2, with 5 days and 25 days combined and FDR < 0.05. (C) Representative confocal images at the level of cell-bodies (top) and mid-lamina (bottom) of C2/3 neurons co-expressing CD8::GFP and smGdP, α-synuclein[A53T], tau[P301L] as well as mini-w+ control. Arrow heads indicate dystrophic neuronal processes (top) and the anatomic location of C2 synaptic varicosities (bottom) at the lamina surface (Tuthill et al., 2013). (D) Quantification of optic lobes with dystrophic C2/3 neuronal processes. Fractions are shown. N = number of optic lobes, noted at bottom of bars. (E) Quantification of C2 CD8::GFP lamina projection width. Replicate values of different optic lobes, mean and SD are shown. ns, p > 0.05; **p < 0.01; ***p < 0.001; one-way ANOVA with Dunnett’s multiple comparison test.

To validate whether the magnitude of transcriptional deregulation identifies preferentially vulnerable neurons with a neurodegeneration-relevant read-out, we expressed mutant tau or α-synuclein in C2 and C3 neurons together with a membrane-tagged GFP (CD8::GFP) in order to evaluate neuronal structural demise. C2 and C3 neurons are an interesting test case, since they are significantly more affected in P301L tau *versus* A53T α-synuclein brains (Figure S6G-S6I), which is particularly apparent at 25 days of age (Figure S5B and S5D). Their cell bodies of are located in the optic lobes, which send projections into the laminae (Tuthill et al., 2013). In 25 day old P301L tau animals C2/3 neurons show dystrophic neuronal processes and >50% thinning of the terminal GFP-labeled varicosities in comparison to animals expressing smGdP or to mini-*w*^*+*^ controls (Figure 3C-3E). Conversely, animals that express A53T α-synuclein in C2/3 neurons do not show morphological defects (Figure 3C-3E). Hence, our analyses, based on the number of DEGs, is capable of identifying resilience/vulnerability to pathogenic α-synuclein or tau expression.

### Prediction of vulnerability signatures in human brain cell types

We then used the classification of *Drosophila* neuron types into preferentially vulnerable and resilient to uncover genes and pathways associated with this dichotomy. For both α-synuclein and tau separately, we pooled the RNA expression data of all vulnerable cell types and of all resilient cell types in the mini-*w*^*+*^ control dataset. We then performed differential gene expression testing between vulnerable and resilient cells to uncover predictive signatures to A53T α-synuclein and to P301L tau-induced stress (Figure 4A and Supplemental file 1). We then performed gene ontology (GO) term overrepresentation analysis on the resulting differentially expressed genes (Figure 4B and 4C, Figure S7A and S7B). This identifies processes related to mitochondrial energy metabolism expressed at higher levels in neuron cell types that are resilient in A53T α-synuclein expressing animals (Figure 4B), suggesting that high baseline mitochondrial respiration capacity might be protective to α-synuclein. This finding reinforces an established link between α-synuclein and mitochondrial dysfunction (Devi et al., 2008; Ganjam et al., 2019). P301L tau resilient neurons exhibit higher expression of ‘axon guidance’ and ‘synapse organization’ terms (Figure 4C), supporting the idea that genes involved in the development or remodeling of these compartments might be able counteract tau-induced synaptic loss. GO terms enriched *both* in A53T α-synuclein and P301L tau vulnerable neurons are related to translation, dendrite, synapse and axon (Figure S7A and S7B), suggesting that distance from the neuronal soma as well as increased input into the protein quality control network are vulnerability factors. Also ion channel and Ca^2+^ related terms are enriched in vulnerability genes, consistent with a role of neuronal excitability and altered Ca^2+^ homeostasis in Parkinson’s and Alzheimer’s disease vulnerable neurons (Fu et al., 2018; Saxena and Caroni, 2011; Sulzer and Surmeier, 2013). Thus, we have uncovered gene expression signatures that associate with preferential vulnerability and resilience to pathogenic α-synuclein and tau by screening >200 fly neuron subtypes.

**Figure 4.**
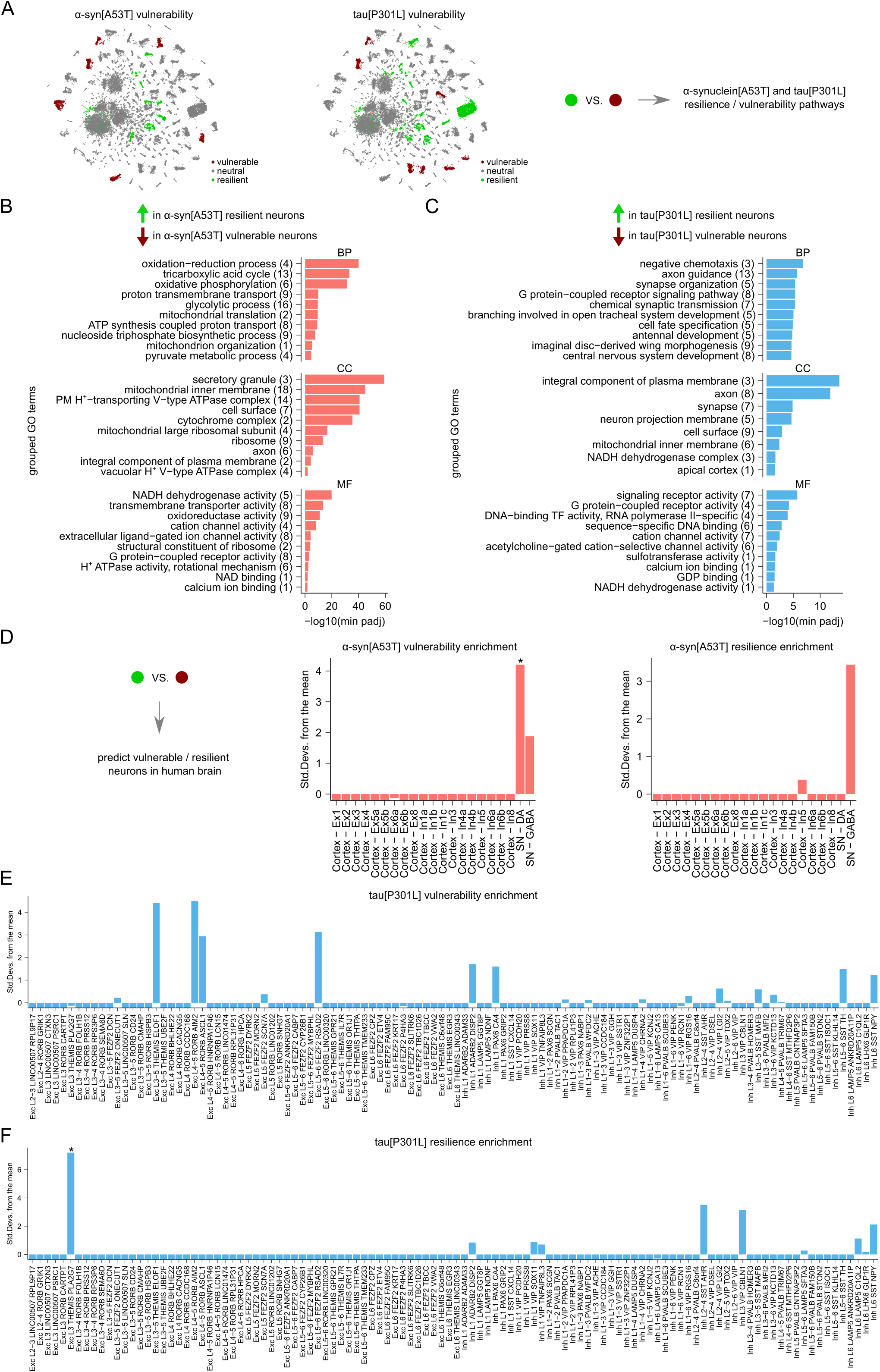
α-synucleinopathy and tauopathy resilience and vulnerability signatures. (A) UMAP projection of mini-w+ control cells where neurons classified as vulnerable/resilient to A53T α-synuclein or P301L tau are highlighted. These subpopulations of cells are compared to obtain differentially expressed genes defining vulnerability/resilience pathways. (B) Grouped GO terms enriched in A53T α-synuclein resilient neurons as compared to vulnerable ones. Number of GO terms grouped under each parent term are shown in brackets. Top 10 parent terms per ontology are shown. BP, biological process; CC, cellular component; MF, molecular function. (C) Grouped GO terms enriched in P301L α-synuclein resilient neurons as compared to vulnerable ones. Number of GO terms grouped under each parent term are shown in brackets. Top 10 parent terms per ontology are shown. BP, biological process; CC, cellular component; MF, molecular function. (D) Expression weighted celltype enrichment (Skene and Grant, 2016) was used to evaluate increased expression of *Drosophila* A53T α-synuclein vulnerability or resilience genes in human *substantia nigra* (SN) and cortex from the middle frontal gyrus (Agarwal et al., 2020). *p < 0.05, Bonferroni correction was applied to bootstrapped p-values from random lists of genes. (E, F) Expression weighted celltype enrichment (Skene and Grant, 2016) was used to evaluate increased expression of *Drosophila* P301L tau vulnerability or resilience genes in different neuron types across multiple human cortical regions from the Allen Brain Map (Allen Brain Map, 2021). *p < 0.05, Bonferroni correction was applied to bootstrapped p-values from random lists of genes.

Next, we used these signatures to assess whether specific human neuron types have enriched expression levels of α-synuclein or tau vulnerability or resilience genes. Since human genes show a high degree of conservation in *Drosophila* (Adams et al., 2000), we ‘converted’ our deregulated fly genes to their human homologues and used expression weighted cell type enrichment in conjunction with single-nucleus RNA sequencing data from healthy human control brains (Skene and Grant, 2016). Across all neurons in a combined dataset encompassing *substantia nigra* and cortex from the middle frontal gyrus (Agarwal et al., 2020) we find the highest ranking A53T α-synuclein vulnerability genes significantly enriched in *substantia nigra* dopaminergic neurons (SD from mean = 4.2, Figure 4D). Conversely, A53T α-synuclein resilience genes are enriched in *substantia nigra* GABAergic neurons (SD from mean = 3.4, Figure 4D). These findings are in line with human neuropathology of PD brains, where α-synuclein aggregates and neuronal loss are key hallmarks of *substantia nigra pars compacta* dopaminergic neurons, but not of the neighboring GABAergic neurons of the *substantia nigra pars reticulata* (Braak et al., 2003; Damier et al., 1999; Gibb and Lees, 1991; Patt et al., 1991).

To then predict neuron-type specific vulnerability to pathogenic tau across the human cortex, we mapped our P301L tau *Drosophila* vulnerability genes to shared neurons across multiple cortical areas from the Allen Brain Map (Allen Brain Map, 2021) (Figure 4E and 4F). We note that *RORB+ AIM2+* and *RORB+ ASCL1+* excitatory neuron types exhibit increased expression of vulnerability genes (SD from mean = 4.5 and 2.9, Figure 4E), in line with previous results identifying *RORB+* excitatory neurons selectively vulnerable in Alzheimer’s disease (Leng et al., 2021). Conversely, excitatory L3-5 *THEMIS+ PLA2G7+* neurons exhibit significant enrichment of resilience genes (SD from mean = 7.2, Figure 4F), suggesting that this neuron type is protected when facing pathogenic tau. Thus, by comparing vulnerable and resilient neurons in *Drosophila* models of α-synuclein and tauopathy we uncover disease-linked vulnerability and resilience signatures that enable us to prioritize neuron-types in the human brain with altered vulnerability-resilience equilibrium.

## Discussion

In this work we dissect mutant α-synuclein and tau as primary causes of neurodegeneration, establish neuronal identity as a modifier of the associated toxicity, provide gene signatures of vulnerability/resilience to pathogenic α-synuclein and tau and thus pave the way to leverage the brain’s own resilience strategies to uncover new treatments for α-synuclein and tauopathies.

α-synuclein and tauopathies are characterized by differential neuronal vulnerability, resulting in the disease-defining clinical symptoms (Fu et al., 2018). No clear correlation of α-synuclein and tau expression levels with neuronal vulnerability in mutation carriers was apparent in recent single nucleus RNA sequencing datasets from multiple human brain areas. This suggests that neuronal identity is capable of modulating the toxicity of pathogenic α-synuclein and tau. We experimentally confirmed this notion by deep phenotyping and brain-wide single-cell RNA sequencing of novel *Drosophila* models of α-synuclein and tauopathy, where we observe selective neuronal vulnerability independent and beyond the contributions of transgene levels.

Our study also enables the direct comparison of cell-type specific vulnerability/resilience arising from α-synuclein with that elicited by tau, since all transgenes are inserted in the same genomic locus and thus not subject to differential silencing (Feng et al., 2000). Indeed, while cell-type specific α-synuclein and tau levels were highly correlated across all brain cell types in these models, we noted the impairment of different neuronal circuits between the two transgenes. *How* pathogenic α-synuclein and tau affect *different* cell types is an open question, especially since it is thought that both proteins exert their toxicity through amyloidogenesis (Fitzpatrick et al., 2017; Guerrero-Ferreira et al., 2018; Li et al., 2018; Shi et al., 2021; Sun et al., 2020). We also note neuron subclasses in our *Drosophila* models that are vulnerable/resilient to α-synuclein *and* tau proteopathic stress, as well as shared deregulated genes and pathways. This is in line with a partial overlap of the associated human diseases and frequent co-occurrence of α-synuclein and tau aggregates in the same brain regions (Moussaud et al., 2014; Yan et al., 2020). It will be interesting to include other amyloidogenic toxic proteins that are associated with the preferential degeneration of very different neuron types, such as polyglutamine expanded huntingtin (striatal medium spiny neurons), different ataxins with polyglutmaine expansions (cerebellar Purkinje cells) as well as mutant TDP-43 (upper/lower motor neurons) (DiFiglia et al., 1997; Holmberg et al., 1998; Neumann et al., 2006).

We provide proof of concept that it is possible to use the >200 brain neuron types of the fly brain as virtual reaction tubes to ask what cellular signatures correlate with vulnerability and resilience to pathogenic toxic proteins. The proposed pathways associated with α-synuclein and tau vulnerability/resilience – such as mitochondrial, axogenesis and ion channel related mechanisms - markedly overlap with those known in sporadic α-synuclein and tauopathies (Fu et al., 2018; Leng et al., 2021; Roussarie et al., 2020; Sulzer and Surmeier, 2013). This validates our approach and reinforces the causality of pathogenic α-synuclein and tau in the more common sporadic diseases. Since we obtain these results by comparing multiple vulnerable with resilient neuron types, our analysis is focused on shared genes and pathways and thus eliminates many ‘silent bystander marker genes’. Nevertheless, individual or groups of genes will ultimately have to be experimentally tested for their ability to modulate pathogenic α-synuclein or tau toxicity to establish causality and obtain potential new drug targets. In Parkinson’s disease, *substantia nigra* dopaminergic neurons accumulate α-synuclein aggregates, degenerate and cause the disease-defining motor symptoms. We detect this neuron type to be significantly enriched for α-synuclein vulnerability genes, suggesting that its molecular identity predisposes it for α-synuclein mediated toxicity. In addition, high baseline α-synuclein expression levels together with the enrichment of Parkinson’s disease Mendelian and sporadic disease genes seem to further contribute to the selective vulnerability of this cell type (Kamath et al., 2022). Interestingly, we found α-synuclein expression to be relatively low in GABAergic as compared to excitatory neurons. This finding could explain why GABAergic neurons do not typically exhibit α-synuclein aggregates (Surmeier et al., 2017; Wakabayashi et al., 1995). Moreover, *substantia nigra* pars reticulata GABAergic neurons were enriched for *Drosophila* pathogenic α-synuclein resilience genes, providing a possible additional explanation why this cell type is relatively spared in Parkinson’s disease (Braak et al., 2003; Damier et al., 1999; Gibb and Lees, 1991; Patt et al., 1991).

Our models only capture specific forms of α-synuclein and tauopathy, thus leaving other interesting aspects related to cell-type specific vulnerability unexplored. We selected the A53T α-synuclein and P301L tau mutations due to their increased aggregation propensity (Flagmeier et al., 2016; Nacharaju et al., 1999; Strang et al., 2018) and therefore do not capture neuron-type specific vulnerability related to other mutations, which can result in different phenotypes (Schneider and Alcalay, 2017; Spillantini and Goedert, 2013). We have also not explored neuron type specific changes in alternative splicing into three-or four-repeat tau. Finally, our analysis has considered cell-autonomous effects of different neuron types but not multi-cellular effects such as those resulting from neuronal connectivity and spreading of misfolded α-synuclein and tau species (Clavaguera et al., 2009; Courte et al., 2020; Henderson et al., 2019). These might act as an additional layer of complexity on top of a neuron’s intrinsic vulnerability to pathogenic α-synuclein and tau.

## Materials and Methods

### *Drosophila* models

To create new *Drosophila* models for this study, we first generated pQUASTattB by transferring the ‘5xQUAS-hsp70 promoter-MCS-SV40 polyA signal’ cassette from pQUAST (Addgene #24349, (Potter et al., 2010)) into pUASTattB (Bischof et al., 2013) via BamHI digestion and T4 ligation (NEB). We linearized pQUASTattB or pUASTattB with EcoRI-XhoI to enable the insertion of transgenes via Gibson assembly (HiFi DNA Assembly, NEB). V5-tagged, fluorophore-dead GFP (smGdP::V5; to enable downstream experiments with GFP based reporters together with this line) was amplified from pJFRC206 (with primers smGFP::v5_hifi_F/R from Addgene #63168, (Nern et al., 2015)); human wild-type and A53T mutant α-synuclein coding sequence was custom synthesized (IDT, sequences below) and tau[P301L] 0N4R amplified from previously reported constructs with tau_hifi_F/R (Zhou et al., 2017). The α-synuclein NACore 68-GAVVTGVTAVA-78 and PHF6 tau 306-VQIVYK-311 deletions were introduced with site-directed mutagenesis with asyn_del_NAC_F/R and tau_del_VQIVYK_F/R (von Bergen et al., 2000; Rodriguez et al., 2015). To control for potential effects of the plasmid insertion itself, we generated an expression deficient variant ‘mini-w+’ by removing the ‘5xUAS-hsp70 promoter’ cassette from pUASTattB via HindIII-XhoI digesting, blunting and ligating with the Quick Blunting Kit (NEB) and T4 ligase. All transgenes were inserted into the VK00005 attP landing site (75A10, 3L: 17,952,108..17,952,108 [-]) by BestGene and backcrossed for five generations into an inbred Canton-S strain harboring the w[1118] mutation (Pech et al., unpublished). nSyb-QF2w.attP2, nSyb-QF2.attP2 and UAS-mCD8::GFP.su(Hw)attP1 where obtained from Bloomington Drosophila Stock Centre (51960, 78338, 32187); the C2.C3_d1-Gal4 line (R20C11-p65ADZp_attP40; R48D11-ZpGDBD_attP2) from Janelia FlyLight (SS00779). Flies were maintained on standard corn meal and molasses food. For all experiments, male flies of control and experimental genotypes were raised in parallel on the same batch of food in a temperature and light-controlled incubator at 25°C and 12-hour light-dark cycle.

smGFP::v5_hifi_F CTCTGAATAGGGAATTGGGCAAACATGGGAAAACCTATACCGA

smGFP::v5_hifi_R AAAGATCCTCTAGAGGTACCCTTAGGTACTATCCAGTCCC

tau_hifi_F AACTCTGAATAGGGAATTGGGCAAACATGGCTGAGCCCCGCCAGG

tau_hifi_R AAAGATCCTCTAGAGGTACCCTTACAAACCCTGCTTGGCC

asyn_del_NAC_F CAAAAGACCGTCGAGGGA

asyn_del_NAC_R TCCTACGTTGGTTACCTG

tau_del_VQIVYK_F CCAGTTGACCTGAGCAAGGTGACC

tau_del_VQIVYK_R ACTGCCGCCTCCCGGGAC

α-synuclein sequence: CTCTGAATAGGGAATTGGGCAAACATGGACGTCTTTATGAAGGGACTGAGTAAAGCGAAGGAGGGTGTGGTGGCGGCCGCAGAAAAAACGAAGCAGGGCGTGGCAGAGGCGGCGGGAAAGACGAAGGAAGGAGTGCTGTATGTTGGCTCGAAAACCAAGGAAGGCGTCGTGCACGGCGTTGCCACAGTTGCGGAGAAGACGAAGGAGCAGGTAACCAACGTAGGAGGAGCTGTAGTAACAGGCGTAACCGCCGTTGCCCAAAAGACCGTCGAGGGAGCTGGTTCGATAGCCGCTGCTACCGGTTTTGTAAAAAAAGATCAGTTGGGGAAAAACGAAGAGGGTGCTCCGCAGGAGGGAATCCTGGAAGACATGCCGGTGGACCCAGATAATGAGGCATACGAGATGCCATCCGAAGAAGGCTACCAGGATTATGAACCAGAGGCATAAAGGATCTTTGTGAAGGAAC

α-synuclein[A53T] sequence: AACTCTGAATAGGGAATTGGGCAAACATGGACGTCTTTATGAAGGGACTGAGTAAAGCGAAGGAGGGTGTGGTGGCGGCCGCAGAAAAAACGAAGCAGGGCGTGGCAGAGGCGGCGGGAAAGACGAAGGAAGGAGTGCTGTATGTTGGCTCGAAAACCAAGGAAGGCGTCGTGCACGGCGTTACCACAGTTGCGGAGAAGACGAAGGAGCAGGTAACCAACGTAGGAGGAGCTGTAGTAACAGGCGTAACCGCCGTTGCCCAAAAGACCGTCGAGGGAGCTGGTTCGATAGCCGCTGCTACCGGTTTTGTAAAAAAAGATCAGTTGGGGAAAAACGAAGAGGGTGCTCCGCAGGAGGGAATCCTGGAAGACATGCCGGTGGACCCAGATAATGAGGCATACGAGATGCCATCCGAAGAAGGCTACCAGGATTATGAACCAGAGGCATAAAGGATCTTTGTGAAGGAACCT

### Western blots

Fly heads from 5 flies (3 samples collected per genotype) were homogenized using a motorized pestle and 50 μL RIPA buffer (Sigma) containing protease and phosphatase inhibitors (cOmplete and PhosStop, Roche). After incubation on ice for 10 minutes, samples were spun down at 16000 g for 10 minutes, supernatant was collected and protein concentration quantified using the 660 nm Protein Assay Reagent (Pierce) supplemented with 50 mM Ionic Detergent Compatible Reagent (Pierce) in a GloMax Multi Detection Plate Reader (Promega). After boiling in 1X Laemmli Sample Buffer (Bio-Rad) with 2.5% 2-mercapto-ethanol (Sigma), 5 μg of head lysate or 5 μg of Human Normal Brain GenLysate (G-Biosciences) was subjected to standard SDS-PAGE using a NuPage 4-12% Bis-Tris gel (Thermo Fisher Scientific) and transferred onto a nitrocellulose membrane (BioRad) using the Trans-Blot Turbo Transfer System (Bio-Rad). Subsequently, total tranferred protein levels were quantified using Ponceau S (Sigma) or the No-Stain Labeling Reagent (Thermo Fisher Scientific) with an iBright imaging system (Thermo Fisher Scientific). Membranes were washed in tris buffered saline (TBS), blocked for 1 h in 3% BSA in TBS with 0.05% Tween-20 (TBS-T) and incubated overnight at 4°C with primary antibodies in 1% BSA in TBS-T. The following day, membranes were washed and incubated with HRP-conjugated secondary antibodies for 1.5 h at room temperature in 5% milk powder in TBS-T. After washing, HRP signal was detected using the Western Lightning Plus-ECL, Enhanced Chemiluminescence Substrate (Perkin-Elmer) with the iBright imaging system and quantified with Fiji. The following antibodies were used: Mouse anti-tau (1:1000, HT7, Thermo Fisher Scientific), mouse anti-α-Synuclein (1:1000, clone 42, BD Biosciences) and an HRP-conjugated secondary anti-mouse antibody (1:5000, Jackson ImmunoResearch).

### Electroretinograms

Electroretinograms (ERGs) were recorded as previously described (Slabbaert et al., 2016). Briefly, flies were immobilized on glass slides with 5 s fix UV glue (JML). Glass electrodes (borosilicate, 1.5 mm outer diameter, Hilgenberg) were filled with 3 M NaCl. One electrode was positioned in the thorax as a reference while another was placed on the fly eye for recording. Responses to repetitive light stimuli were measured using Axosope 10.7 (Molecular Devices) and analyzed with Clampfit 10.7 (Molecular Devices) and Igor Pro 6.37 (WaveMetrics).

### Immuno-histochemistry

Immunohistochemistry of aged adult brains was performed by dissecting brains in ice-cold PBS and fixing them with 3.7% formaldehyde in PBS with 0.3% Triton X-100 (PBX) for 20 min at room temperature. Subsequently, brains were washed in PBX, blocked with 5% goat serum in PBX and overnight incubated with primary antibodies in 5% goat serum in PBX at 4°C. Subsequently, samples were washed and incubated with secondary antibodies in 5% goat serum in PBX overnight at 4°C. After final washes brains were mounted in RapiClear 1.47 (SunJin Lab). The following antibodies were used: Rabbit anti-GFP (1:500, A11122, Thermo Fisher Scientific), mouse anti-Brp (1:100, nc82, DSHB), Alexa Fluor 488 conjugated anti-rabbit (1:1000, Thermo Fisher Scientific) and Alexa Fluor 555 conjugated anti-mouse (1:1000, Thermo Fisher Scientific). Stained brains were imaged on a Nikon A1R confocal microscope and a Nikon Plan-Apochromat 20x 0.75 NA objective. Z-stacks of optic lobes and attached laminae were acquired encompassing C2 and C3 neuronal cell-bodies and their projections. Before analysis, image names were blinded by assigning random numbers. Subsequently, mid-level slices through cell-body layer or laminae were extracted and the number of images with dystrophic neurons adjacent to the cell body layer was classified and C2 neuronal terminal projection width measured in Fiji.

### Brain sections

Heads of 5, 25 and 45 day old flies were immediately fixed after decapitation in 4% paraformaldehyde (Laborimpex) and 2% glutaraldehyde (Polysciences, Inc) in 0.1 M Na-Cacodylate buffer pH 7.4 (Merck) for 2 h at room temperature. Samples were further fixed at 4°C overnight, washed with 0.1 M Na-Cacodylate pH 7.4, and subsequently osmicated with 2% osmium tetroxide (Laborimpex). After staining in 2% uranyl acetate solution (EMS) for at least 1.5 h and dehydration in an ascending series of ethanol solutions, samples were embedded in Agar 100 (Laborimpex) and cured at 60°C for 48 h. Alternatively, the samples were kept substantially longer in 2% osmium tetroxide and kept overnight in 0.5% uranyl acetate/25% methanol solution before continuing with the dehydration and embedding steps. Semi-thin sections (1.5 μm) of the fly heads were collected on microscopy slides. The sections were then dried and stained on a heating block with a 1% toluidine blue (Merck) solution including 2% Borax for 90 s at 60°C. The stained sections were mounted with Eukit Quick-hardening mounting medium (Sigma) and imaged at the level of the fan-shaped body with a Leica DM25000M microscope using a 20x HC PL Fluotar Ph2 Air NA 0.50 objective and the LAS V4.0 software. Vacuoles in the fly brain were automatically extracted in the thresholded image using the analyze particles tool (with parameters: size = 1.767-Infinity, which is equivalent to a minimum 1.5 μm vacuole diameter, and circularity = 0.40-1.00) in ImageJ and the total vacuole area per brain area calculated.

### Fly sleep and activity monitoring

Profiling of sleep and activity behaviour of 5, 25 and 45 day old male flies was performed using the fly behaviour video recording platform ethoscope (Geissmann et al., 2017). Briefly, flies were sorted into glass tubes (65 mm length, 5 mm external and 3 mm internal diameter) with standard corn meal and molasses food on one end and closed with cotton wool on the other. 20 individual animals per ethoscope were recorded for at least 4 consecutive days at 2 frames per second with infrared light in incubators at 25°C with 12 hour light-dark conditions. The position of each animal was saved at each time point in SQLite files and subsequently analysed with R (v 3.6.3) and rethomics with adjusted R packages behavr, scopr and sleepr (v0.3.99). The behaviour of individual flies was determined with the automatic behaviour annotator at a resolution of 10 s. Thus, in 10 s intervals flies were scored as moving or asleep. Sleep was defined following the 5-min rule, in which immobility bouts longer than 5 min were counted as sleep bouts, including the first 5 min. A fly was scored as immobile if the distance between 2 consecutive frames in the 10 s interval was less than 0.27 mm. Flies that died during the monitoring as well as the first day of recording due to habituation were excluded from further analysis. To describe the activity and sleep profile of individual flies the following parameters were analysed: Within 3 days of the indicated age, the sleep amount per light condition (sleep_fraction_day) and per dark condition (sleep_fraction_night) were calculated. For the latency parameter, the time between zeitgeber 12 (= lights off) and the first sleep bout was determined. The mean bout length as well as the number of sleep bouts were scored during the dark condition. Morning and evening anticipation were calculated as described previously (Valadas et al., 2018). The velocity_if_awake was assessed based on 10 s interval annotation ‘moving’. The cumulative distance within the 10 s interval was calculated and summed per 24 h to determine the travelled distance (total_distance) for each animal. Finally, the means of the activity and sleep parameters were calculated for each genotype, scaled and hierarchically clustered based on correlation distance and average-linkage with the R pheatmap package.

### Single-cell dissociation

Three cohorts of male nSyb-QF2w > mini-w+, nSyb-QF2w > QUAS-smGdP::V5, nSyb-QF2w > QUAS-α-synuclein[A53T] and nSyb-QF2w > QUAS-tau[P301L] were aged in parallel to 5 or 25 +/-1 day of age at 25°C in a 12-hour light-dark cycle. From each genotype in these six repeats, five brains with laminae largely removed were dissected in PBS, which was supplemented with 5 μM Actinomycin D (Sigma) to block changes in gene expression during sample preparation (Wu et al., 2017). To avoid confounding of genotype with experimental batch, all four genotypes were dissected and processed in parallel. To reduce variability between the different experimental batches, reagents and conditions were kept constant. Dissections were carried out directly after the fly incubator light was switched on (ZT0), completed in less than 1 hour by the same experimenters and the genotype-assignment to the dissectors as well as dissection order was alternated. Fly brains were dissociated in 250 μl of 0.6 mg/ml Dispase I (Sigma), 30 mg/ml Collagenase I (Thermo Fisher Scientific), 0.5x Trypsin (Thermo Fisher Scientific) and 5 μM Actinomycin D for 20 min at 25°C 1000 rpm with four triturations in 5 min intervals. Cells were washed once in PBS + 5 μM Actinomycin D and filtered through a 10 μm strainer (Pluriselect) in 60 μl PBS + 0.04% BSA (Sigma).

### Single-cell capture and library preparation

Cells were counted with the FLUNA-FL Dual Fluorescence Cell Counter and viability determined with Acridine Orange/Propidium Iodide dyes (Logos biosystems), which was >93% for all samples, typically 98-100%. We targeted 10 000 captured cells per genotype with Chromium v3.1 chemistry (10x Genomics). All genotypes were run in parallel on multiple lanes of the same Next GEM chip. In different repeat experiments the assignment of genotypes to lanes was alternated. After single-cell encapsulation, library preparation followed the manufacturer’s protocol (Single cell 3’ reagent kits v3.1 user guide; CG000204 Rev D). Briefly, reverse transcription was performed on a C1000 Touch Thermal Cycler (Bio Rad) at 53°C 45 min and 85°C 5 min with a 4°C hold, followed by breakage of the single-cell emulsion, Dynabeads MyOne SILAN clean-up and cDNA amplification at 98°C 3 min, 13 cycles of 98°C 3 min - 63°C 20 s - 72°C 1 min, with a final 72°C 1 min extension and 4°C hold. Amplified cDNA was cleaned with SPRIselect (Beckman Coulter). Thereafter cDNA was enzymatically fragmented, followed by end-repair, A-tailing, SPRIselect clean-up, adapter ligation and another SPRIselect clean-up was carried out prior to PCR amplification of the ligation products incorporating unique sample indices at 98°C 45 s, 15 cycles 98°C 20 s - 54°C 30 s - 72°C 20 s - with a final 72°C 1 min extension and 4°C hold. SPRIselect clean-up of the amplified libraries was carried out. At different check points the quality of cDNA and final sequencing library was evaluated using Qubit (ThermoFisher) and Bioanalyzer (Agilent). Next-generation sequencing (VIB Nucleomics Core) of the single-cell libraries was carried out on a NovaSeq 6000 (Illumina) platform with the following read configuration: Read 1 28 cycles, i7 Index 8 cycles and Read 2 91 cycles. Libraries were first sequenced shallow to determine library complexity and subsequently sequenced deeper to achieve ∼80% sequencing saturation.

### Data pre-processing

Pooled sequencing data was demultiplexed and index-hopping-filter v1.0.1 (10x Genomics) applied in order to minimize misassignment of reads to wrong samples. This resulted in the exclusion of 1-2% of the acquired reads. Subsequently the feature-barcode matrix was computed with the Cell Ranger count command (v4.0.0, 10x Genomics) with the –expect-cells flag set to our targeted 10 000 cell recovery. As a reference for mapping the 4^th^ 2020 FlyBase *Drosophila melanogaster* release was used (r6.35). In order to also detect transgenes, their coding sequence plus SV40 terminator sequences were added as artificial chromosomes. The fraction of contaminating ambient RNA was automatically determined and corrected with the soupX (v1.5.2) autoEstCont() and adjustCounts() commands using default parameters. Ambient RNA was estimated to range between 2 and 10% in our samples. Subsequently, individual ambient RNA corrected matrices were loaded into scanpy (v1.8.1) and concatenated into one dataset. Quality control steps included filtering of cells from 5 day old animals at a minimum of 800, 25 day old animals 600 detected genes as well as a maximum of 20 000 counts and 15% mitochondrial content. We chose a lower gene detection threshold in older animals since it is known that the RNA content of *Drosophila* brain cells decays exponentially with age (Davie et al., 2018).

### Clustering and cell-type identification

For downstream visual exploration, clustering and cell-type identification, soupX corrected UMIs were normalized to 10 000 per cell and log1p-transformed. The top 2000 most highly variable genes were determined with the scanpy implementation of the original Seurat approach, with transgenes excluded (Satija et al., 2015). These highly variable genes were then used to reduce the dimensionality of the data-set with PCA. We explored 30, 50, 70, 90 and 110 PCs and the resulting separation of pretrained cell-type classifier predictions (see below) on a 2-dimensional UMAP projection. This led to the choice of 70 PCs. Similarly, for Leiden based clustering we explored several resolutions qualitatively and chose 3 as a comparably low base resolution, to avoid oversplitting into small sub-cell types and resulting loss of statistical power in differential gene expression testing. To link our Leiden clusters to cell types, we integrated the predictions of two previously trained fly brain cell type classifiers (Pech et al., unpublished; Özel et al., 2021). Each cell in our dataset was assigned a cell-type label with both models and for each cluster the proportions of cell-type predictions were calculated. For the majority of Leiden clusters both models were in agreement for >90% of its cells, in which case we transferred the majority cell-type name to this cluster. In some instances, clusters required further splitting, in which case we increased the Leiden resolution in steps of 1 until we yielded unambiguous single cell-types. In addition, in our cell-type assignments we also leveraged cell-type specific bulk transcriptomes by calculating Pearson correlation coefficients with the sum aggregated highly variable genes in our Leiden clusters (Davis et al., 2020; Konstantinides et al., 2018). Broad cell-type annotations were inferred from classifier derived cell-type labels as well as neuronal (elav, nSyb), glia (repo), photoreceptor (chp) and hemocyte (Hml) marker genes. Neurotransmitter identities were assigned based on previously established neuron-neurotransmitter connections (Davis et al., 2020) as well as Acetylcholine (VAChT, ChAT), Glutamate (VGlut), GABA (Gad1), Dopamine (ple), Tyramine/Octopamine (Tdc2) and Histamine (Hdc) metabolism marker genes.

### Differential gene expression testing and GO analysis

Cell-type specific differential gene expression (DEG) testing of A53T α-synuclein, P301L tau and smGdP vs. non-expressing cells in mini-w+ was carried out in a cell-type specific pseudobulk manner, which has been shown to better control for false discoveries (Crowell et al., 2020; Squair et al., 2021; Zimmerman et al., 2021). Therefore, UMIs in each gene per cell type of each genotype and repeat was sum-aggregated and rounded to the next integer, since soupX correction yields floating point numbers and the below steps expect integer inputs. We then used DESeq2 and edgeR to fit negative binomial models and carry out the Wald- and quasi-likelihood F-tests respectively for differential gene expression testing (Love et al., 2014; McCarthy et al., 2012; Robinson et al., 2009). We removed experimental batch as a covariate according to the design formula ∼date+genotype and performed our comparisons at 5 days, 25 days and 5 plus 25 days combined. All genes below a Benjamini-Hochberg corrected p-value of 0.05 were considered as deregulated. For interpretation of the magnitude of cell type specific transcriptomic changes we only retained cell types that had a minimum of 10 cells/repeat as well as a minimum total of 65 or 130 mutant cells in the age-separated or age-merged analysis. In addition, cell types either had to be identified by name or be clear free-standing clusters (as opposed to the different clusters that are densely packed in the center of the UMAP).

To extract DEGs between neuron types predicted to be vulnerable or resilient (see below), we used a similar approach. We first sum-aggregated UMIs across genes of all vulnerable or resilient neurons in the mini-w+ dataset and rounded the result to the next integer. We then used DESeq2 and the Wald test to compare all resilient with all vulnerable neurons while regressing out experimental date. All genes below a Benjamini-Hochberg corrected p-value of 0.05 were considered as deregulated. For GO term enrichment analysis we used the enrichGO() function from the clusterProfiler (v3.18.1) and org.Dm.eg.db (v3.12.0) R packages (Carlson, 2019; Yu et al., 2012). The minimum gene set size was 10, maximum 500 and background defined by all detected genes in vulnerable and resilient neurons. GO terms with Benjamini-Hochberg adjusted p-values < 0.05 were retained and grouped by parent terms based on similarity and the ‘Wang’ method (Wang et al., 2007) in the mgoSim() function of the GOSemSim R package (v2.16.1) and the reduceSimMatrix() functions in the rrvgo (v1.2.0) package as implemented in the go_reduce() wrapper (Feleke et al., 2021).

### Neuron-type classification

We modelled the number of DEGs resulting from DESeq2 and edgeR differential gene expression testing at 25 days and 5 + 25 days with negative binomial regression in order to identify neurons with larger or lower transcriptional deregulation than would be expected by the neuron-type specific covariates number of cells, number of detected genes and transgene levels. With the log-likelihood ratio test and the Akaike information criterion we validated the negative binomial model as a better fit as compared to the Poisson or intercept only models. Neurons were classified as vulnerable if they had more DEGs than the upper bound of the 95% confidence interval in the DESeq2 and edgeR models at 25 days or 5 + 25 days combined and that were not fewer than the 95% confidence interval in any of these models. Neurons were classified as resilient if they had fewer DEGs than the lower bound of the 95% confidence interval in the DESeq2 and edgeR models at 5 + 25 days combined and did not have more DEGs than the upper bound of the 95% confidence interval in any of the 25 day models.

### Mapping vulnerability-resilience genes to human neuron types

To uncover neuron types in the human brain with increased relative expression of α-synuclein or tau vulnerability/resilience genes as uncovered in *Drosophila* models, we used Expression Weighted Celltype Enrichment (EWCE, v1.0.0) (Skene and Grant, 2016) together with human brain single-nucleus RNA sequencing datasets from *substantia nigra* and middle frontal gyrus (Agarwal et al., 2020) or from multiple cortical areas, including the middle temporal gyrus, anterior cingulate cortex, primary visual cortex, primary motor cortex, primary somatosensory cortex and primary auditory cortex (Allen Brain Map, 2021). *Drosophila* genes up-or downregulated with a log2-fold change >2.5 or <2.5 and a Benjamini-Hochberg corrected p-value <0.05 in vulnerable vs. resilient neurons were mapped to human neuron types by converting them to their human orthologous genes with DIOPT (Hu et al., 2011). In the human datasets, we only retained neurons, since our study focuses specifically on *neuronal* vulnerability. We removed genes with little variance across each of the datasets with the EWCE function drop_uninformative_genes() and calculated the specificity matrix with generate_celltype_data(). To assess whether our gene sets are enriched in any of the human neuron types we used the bootstrap_enrichment_test() function with human orthologous genes of the detected genes in our *Drosophila* dataset as background. This allowed us to compare the average neuron type specific expression of our vulnerability/resilience gene sets with the expression of 10 000 random gene lists of the same length.

## Acknowledgements

We would like to thank Kristofer Davie for feedback and access to the Vlaams Supercomputer Centrum (VSC); Mark Fiers for data analysis advice; Vinoy Vijayan for help; Joost Schymkovitz, Frederic Rousseau, Bart de Strooper, Pierre Vanderhaeghen and Joris de Wit for valuable feedback and members of the Verstreken lab for discussions. We also thank the VIB Nucleomics Core facility, the VIB Bioimaging Core and the VSC for help and the Bloomington Drosophila Stock Center for stocks (NIH P40OD018537). We used data from the Genotype-Tissue Expression (GTEx) Project. GTEx Analysis V8 data was used in this study and obtained on 26/08/2019 from the GTEx Portal. This work was supported by Alzheimer Research Foundation (SAO-FRA 2020/0011 to R.P.) and an ERC consolidator grant (ERC-2014-CoG - 646671), the EU Innovative Medicines Initiative 2 (IMI2), the Chan Zuckerberg Initiative, FWO Vlaanderen (G0A5219N, G0B8119N), Stichting Alzheimer Onderzoek, a Methusalem grant of the Flemish Government, Opening the Future (Leuven University fund) and VIB all to P.V. R.P and N.K. are supported by EMBO Long-Term Fellowships (ALTF 980-2019, ALTF 299-2019), J.J. by a FWO PhD Fellowship (1199518N), E.N. by Interne Fondsen KU Leuven and C.C. by a Marie Skłodowska-Curie Actions - Seal of Excellence FWO Postdoctoral Fellowship. P.V. is an alumnus of the FENS-Kavli Network of Excellence.

## Author contributions

Conceptualization, R.P., P.V.; Methodology, R.P., S.K., N.S., N.K., J.S., E.N., J.J., C.C., P.V; Software, R.P., N.K, J.J., S.K.; Validation, R.P., S.K.; Formal Analysis, R.P., N.K., S.K.; Investigation, R.P., S.K., N.K., E.N., S.P., C.C.; Resources, P.V., S.P., S.A.; Data Curation, R.P., S.K., N.K.; Writing – Original Draft, R.P., P.V., S.K., N.K.; Writing – Review & Editing, R.P., P.V., E.N., S.K., N.K., C.C., S.P., S.A., J.J.; Visualization, R.P., N.K., S.K.; Supervision, P.V., R.P., S.A.; Project Administration, R.P., S.K.; Funding Acquisition, P.V., R.P., N.K., E.N., C.C., J.J., S.A.

## Supplemental figure legends

**Figure S1.**
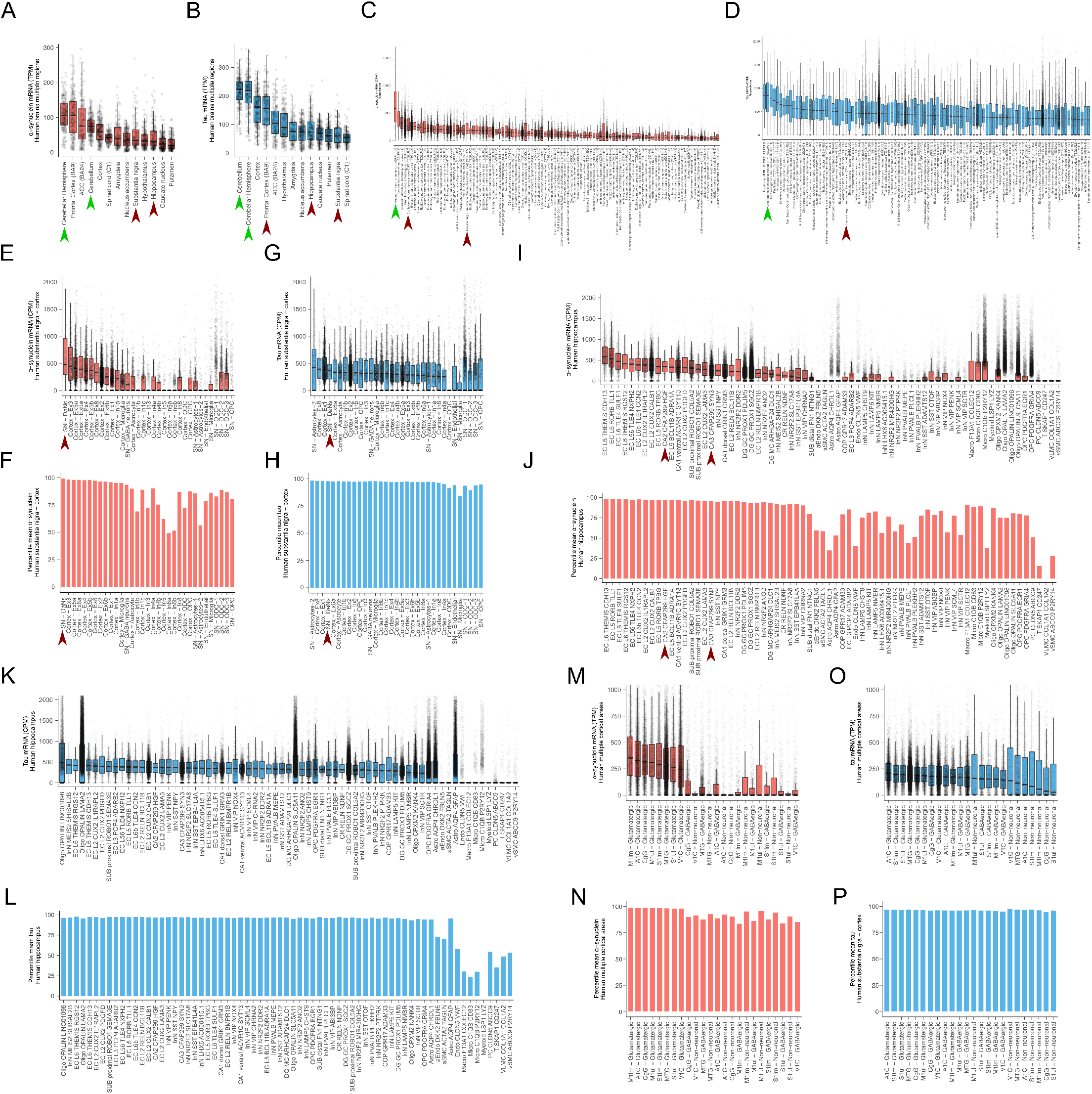
α-synuclein and tau expression levels across multiple regions and cell types of the mouse nervous system and human brains. (A, B) Normalized α-synuclein and tau bulk mRNA expression across 13 different human brain regions from the GTEx consortium. Individual samples, median, interquartile range (IQR) and whiskers at 1.5xIQR are shown. ACC, anterior cingulate cortex. (C, D) Normalized median α-synuclein and tau expression of the top 100 cell types out of 265 from the Zeisel et al. 2018 mouse nervous system atlas is shown. For consistency, only samples processed with Chromium 10x v1 chemistry are shown and cell types with a minimum of 5 cells. Individual cells, median, IQR and whiskers at 1.5xIQR are shown. (E, G) Normalized α-synuclein and tau expression at the cell-type level across human *substantia nigra* and frontal gyrus from the Agarwal et al. 2020 dataset. Individual cells, median, IQR and whiskers at 1.5xIQR are shown. (F, H) Percentile of mean α-synuclein and tau expression amongst the means of genes that are detected in at least 5% of a given cell type in the Agarwal et al. 2020 dataset. SN, *substantia nigra*; DaNs dopaminergic neurons; Ex, excitatory neurons; In, inhibitory neurons, ODC, oligodendrocytes, OPC, oligodendrocyte precursor cell. (I, K) Normalized α-synuclein and tau expression at the cell-type level across human hippocampus, subiculum and entorhinal cortex from the Franjic et al. 2022 dataset. Individual cells, median, IQR and whiskers at 1.5xIQR are shown. (J, L) Percentile of mean α-synuclein and tau expression amongst the means of genes that are detected in at least 5% of a given cell type in the Franjic et al. 2022 dataset. EC, entorhinal cortex; SUB, subiculum; GC, granule cell; MC, mossy cell; Astro, astrocyte; OPC, oligodendrocyte precursor cell; COP, committed OPC; aEndo, arterial endothelial cell; PC, pericyte; vSMC, venous smooth muscle cell; aSMC, arterial smooth muscle cell; VLMC, vascular and leptomeningeal cell. (M, O) Normalized α-synuclein and tau expression at the broad cell-type level across multiple areas from the human cortex from the Allen Brain Map data (Allen Brain Map, 2021). Individual cells, median, IQR and whiskers at 1.5xIQR are shown. (N, P) Percentile of mean α-synuclein and tau expression amongst the means of genes that are detected in at least 5% of a given cell type in the Allen Brain Map data (Allen Brain Map, 2021). MTG, middle temporal gyrus; CgG, anterior cingulate gyrus; V1C, primary visual cortex; A1C, primary auditory cortex; M1ul, upper limb region of primary motor cortex; M1lm, lower limb region of primary motor cortex; S1um, upper limb region of primary somatosensory cortex; S1lm, lower limb region of primary somatosensory cortex. Arrowheads indicate vulnerable (dark red) and not reported as affected (green) example cell types from the text.

**Figure S2.**
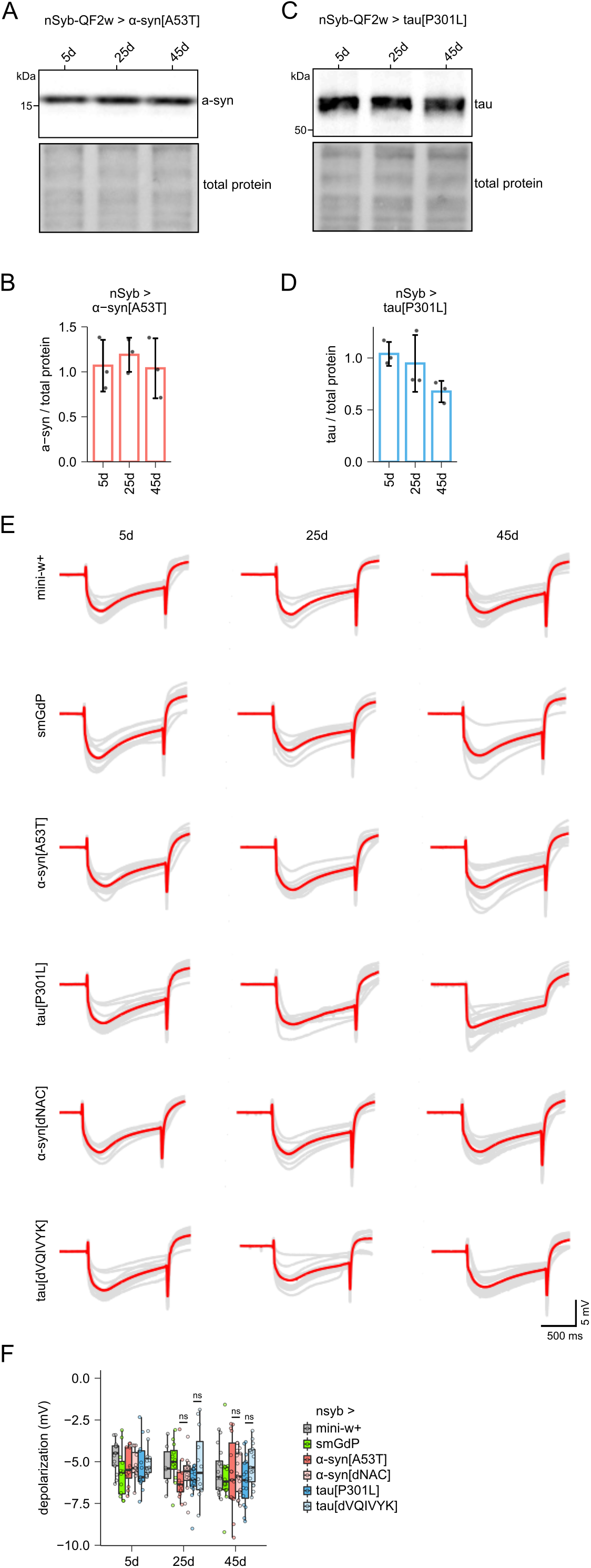
α-synuclein and tau protein levels in ageing fly models, ERG traces and quantification. (A, B) Representative immuno-blot and quantification of α-synuclein protein levels at 5, 25 and 45 days in mutant α-synuclein fly model heads. Replicate values, mean and SD are shown. (C, D) Representative immuno-blot and quantification of tau protein levels at 5, 25 and 45 days in mutant tau fly model heads. Replicate values, mean and SD are shown. (E) ERG traces from individual experiments (gray) as well as their mean (red) are shown. (F) Comparable extend of photoreceptor depolarization is apparent across genotypes and time points. Replicate values, median, IQR and whiskers at 1.5xIQR are shown.

**Figure S3.**
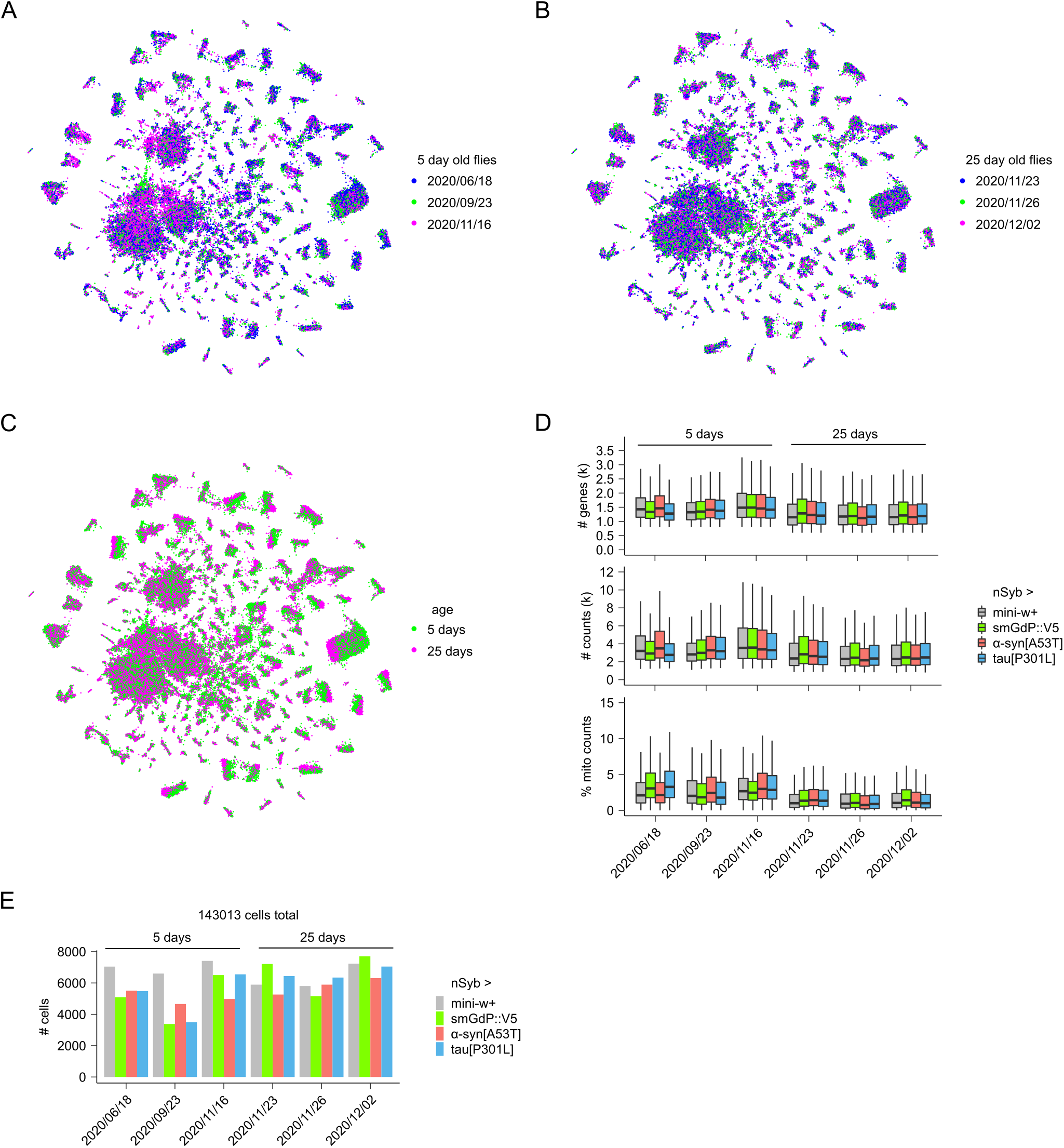
Quality control overview of scRNA-seq across entire α-synuclein and tauopathy model *Drosophila* brains. (A, B) UMAP projection with individual cells colored according to the experimental date. No integration algorithm was used to obtain intermixing of different experimental batches, suggesting that no large batch effects are present. (C) UMAP projection with 5 and 25 day old cells in green and magenta respectively. Incomplete intermixing in some of the clusters is consistent with ageing driving major changes in the cellular transcriptome and in line with previous observations (Davie et al., 2018). (D) Number of genes, counts and percent of reads mapping to mitochondrial genes in quality control filtered cells across the different experiments and genotypes. Median, IQR and whiskers at 1.5xIQR are shown. (E) Total number of retained cells after quality control filtering across the different experiments and genotypes.

**Figure S4.**
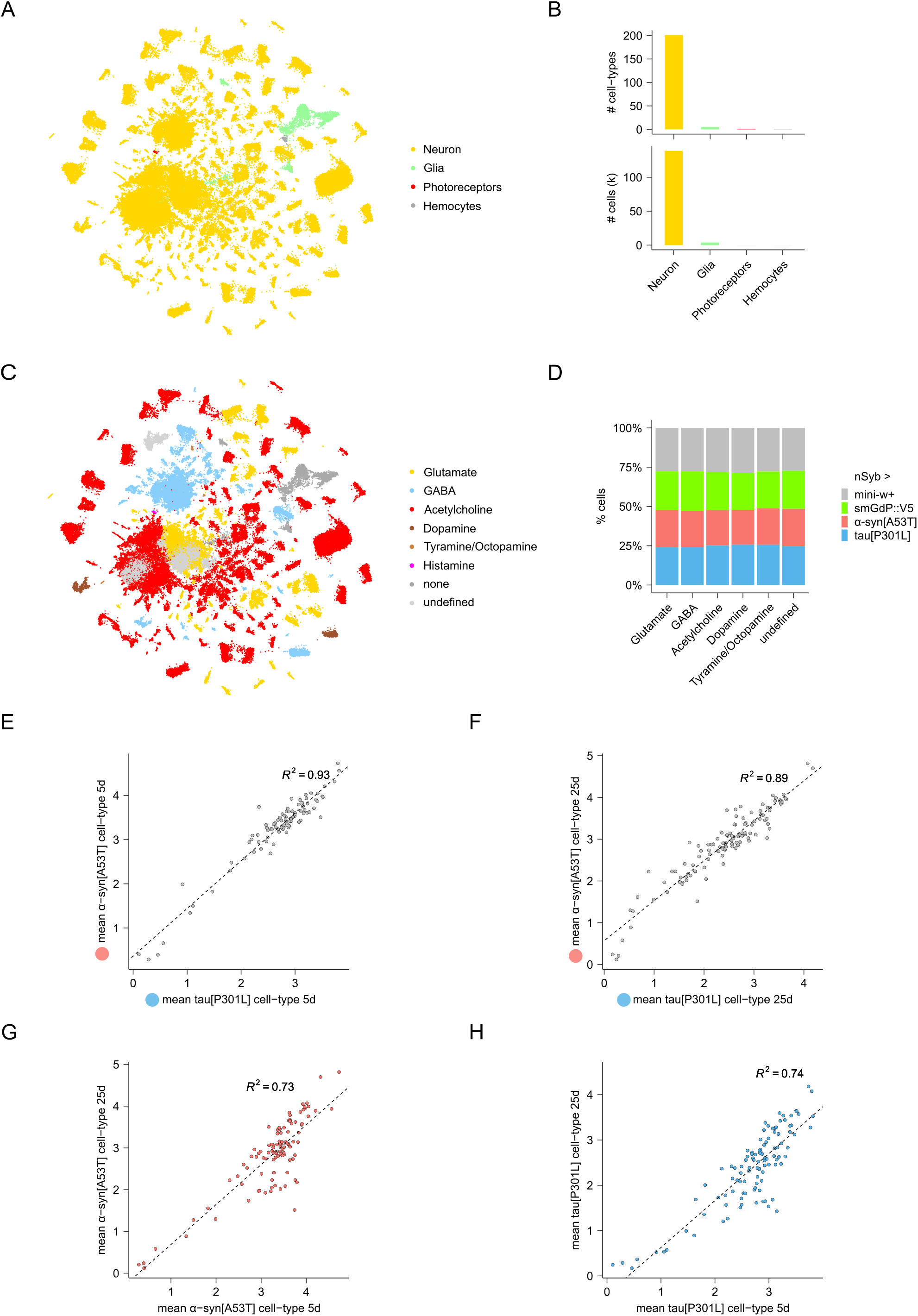
Neuronal diversity detected in scRNA-seq of entire α-synuclein and tauopathy model *Drosophila* brains and α-synuclein and tau expression levels across different cell types and ages. (A) UMAP projection with individual cells colored according to their broad annotation categories. (B) Number of cell types as well as number of individual cells within each broad annotation category. The high neuron to glia ratio is expected in *Drosophila* brains and consistent with previous work (Davie et al., 2018). (C) UMAP projection with individual cells colored according to their main neurotransmitter. Glial cells are labelled as ‘none’. Unidentified neurons 0 and 46 as well as T1 neurons could not be assigned an unambiguous neurotransmitter and were hence labelled as ‘undefined’ (Davis et al., 2020). (D) Percent of cells of each neurotransmitter from each of the three genotypes is shown. (E) Scatter-plot and linear regression of mean normalized and log-transformed transgene levels across cell types of 5 day old P301L tau vs. A53T α-synuclein models. (F) Scatter-plot and linear regression of mean normalized and log-transformed transgene levels across cell types of 25 day old P301L tau vs. A53T α-synuclein models. (G) Scatter-plot and linear regression of mean normalized and log-transformed transgene levels across cell types of 5 vs. 25 day old A53T α-synuclein models. (H) Scatter-plot and linear regression of mean normalized and log-transformed transgene levels across cell types of 5 vs. 25 day old P301L tau models. Throughout (E-H) only cell types with a minimum of 50 cells and those that are not part of the large central clusters are shown.

**Figure S5.**
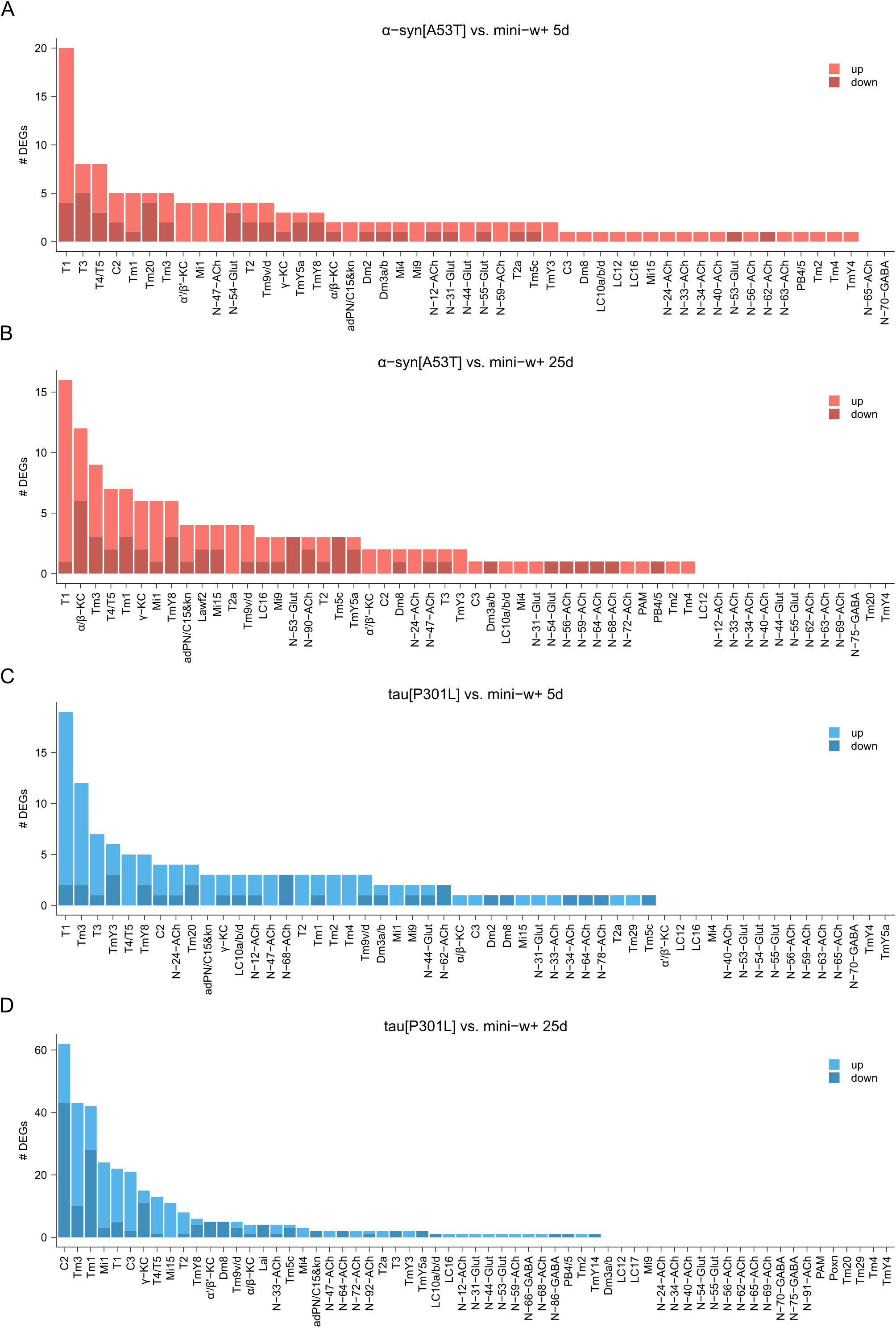
Neuron-type specific transcriptional deregulations in α-synuclein and tauopathy *Drosophila* models. (A-D) The number of significantly up-/downregulated genes per neuron type are shown for A53T α-synuclein and P301L tau vs. mini-w+, calculated for the 5 and 25 day time points separately. Only neuron types that have >10 cells per repeat, >65 total mutant cells and do not map to poorly defined clusters are shown. DEG testing was performed with DESeq2 and FDR < 0.05.

**Figure S6.**
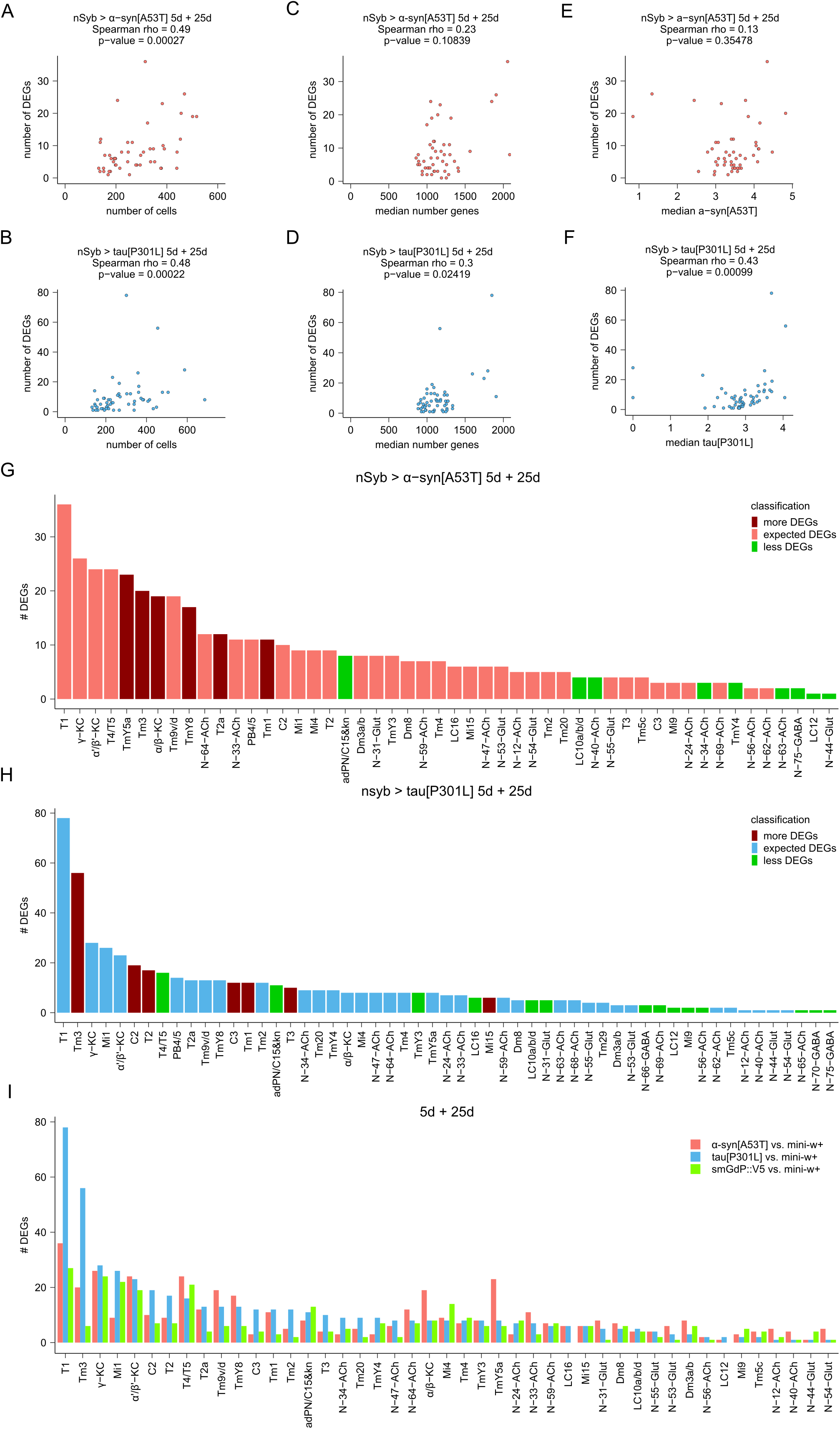
Classification of neuron types based on the magnitude of their transcriptional response. (A-F) Scatter plots and Spearman’s rank correlation coefficients of covariates vs. the number of DEGs obtained with DESeq2, with 5 and 25 days combined and an FDR < 0.05. (A, B) The number of mutant cells, (C, D) the median number of genes and (E, F) transgene levels in a given neuron type are shown. (G, H) The number of DEGs per neuron type are shown for A53T α-synuclein and P301L tau vs. mini-w+ as obtained with DESeq2 at 5 and 25 days of age combined and an FDR < 0.05. Neurons that are transcriptionally more or less deregulated as compared to negative binomial regression models (see Methods) encompassing the above covariates are color coded in dark red or green respectively. (I) The number of DEGs per neuron type are shown for A53T α-synuclein, P301L tau and smGdP vs. mini-w+ as obtained with DESeq2 at 5 and 25 days of age combined and an FDR < 0.05. Only neuron types that have >10 cells per repeat, >130 total mutant cells and do not map to poorly defined clusters are shown.

**Figure S7.**
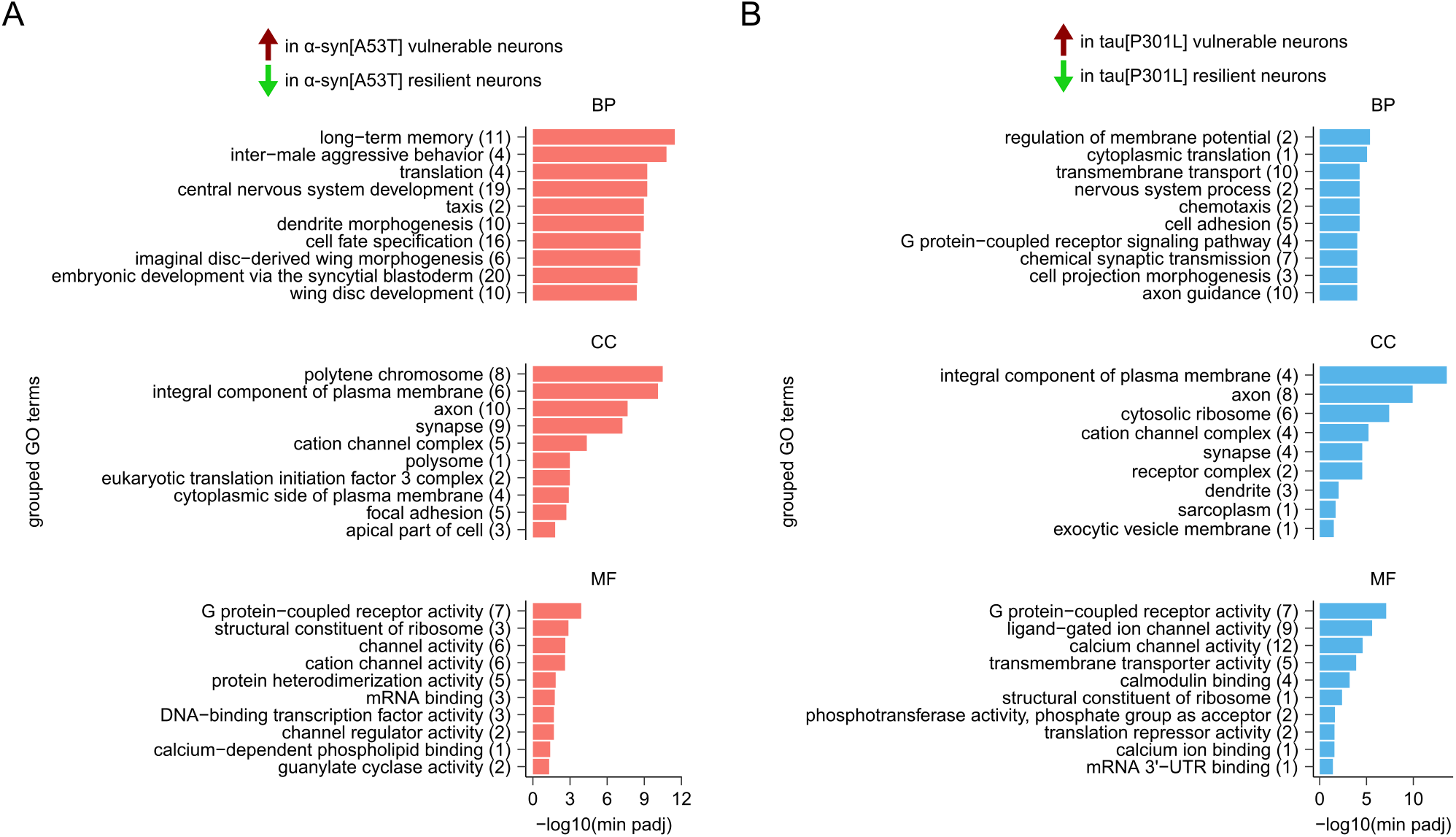
Pathogenic neuronal α-synuclein and tau vulnerability pathways in *Drosophila* brains. (A)Grouped GO terms enriched in A53T α-synuclein vulnerable neurons as compared to resilient ones. Number of GO terms grouped under each parent term are shown in brackets. Top 10 parent terms per ontology are shown. BP, biological process; CC, cellular component; MF, molecular function. (B)Grouped GO terms enriched in P301L tau vulnerable neurons as compared to resilient ones. Number of GO terms grouped under each parent term are shown in brackets. Top 10 parent terms per ontology are shown. BP, biological process; CC, cellular component; MF, molecular function.

**Supplemental file 1. *Drosophila* α-synuclein and tau vulnerability/resilience genes**

A53T α-synuclein and P301L tau vulnerability/resilience signature genes uncovered by comparing all vulnerable with all resilient neuron types in our *Drosophila* models with DESeq2. Top 2000 genes as sorted by Benjamini-Hochberg adjusted p-values.

